# Development of a Synthetic Hydrogel to Foster Microvascularization of an Endometriosis Microphysiological System

**DOI:** 10.1101/2025.10.06.680733

**Authors:** Lauren Pruett, Laura Bahlmann, Ryan Ogi, Angela Jiao, Priyatanu Roy, Matthew Johnson, David Trumper, Linda Griffith

## Abstract

The ascent of novel alternative methods (NAMs) in drug development spotlights the dual needs for improved biological fidelity to *in vivo* along with reproducibility, especially in regulatory applications. The need for pre-clinical models of patient-derived endometriosis lesions motivates development of a vascularizable completely synthetic extracellular matrix (v-CS-ECM) that supports morphogenesis of perfusable microvasculature in a microfluidic device, in the context of relevant lesion cells. This paper describes v-CS-ECM, a peptide-modified polyethylene glycol-based hydrogel crosslinked with a cell-degradable peptide that achieves these dual goals. Vessels form by morphogenesis after the liquid v-CS-ECM precursor, containing endothelial cells and fibroblasts, is injected into the tissue compartment to encapsulate cells. Vessel formation is influenced by ECM biochemical and biophysical properties, source of vascular cells, and microphysiological system (MPS) operating conditions. The v-CS-ECM also supports co-culture of endometrial epithelial organoids (EEOs) and fibroblasts, and formation of microvascularized endometriosis lesion-like structures when all cell types are co-encapsulated in a microfluidic device with constant flow. Hence, v-CS-ECM overcomes limitations of reproducibility and biological function inherent in the fibrin-based ECM typically employed for microvascular morphogenesis, as well as Matrigel for organoid culture, thus offering promise for NAMs evaluating endometriosis drugs in the preclinical setting.

## 1. Introduction

Microphysiological systems (MPS) that capture human physiological processes are emerging as favorable alternatives to animal models in efforts to study disease and develop therapies. Incorporation of perfusable microvasculature is essential for many applications, as capillary endothelia and pericytes not only control the transport of oxygen, metabolites, and therapeutics, they engage in local paracrine signaling of tissue homeostasis and perturbation response, and gate immune cell trafficking^1,2^.

An accessible and now widely-used approach to creating perfusable microvasculature involves morphogenesis of vessels from vascular cells seeded into the tissue chamber of a microfluidic device in a process that embeds them in a supporting extracellular matrix^3–6^ (**Fig. 1**). The morphogenesis process is typically initiated by mixing endothelial cells with supporting cells (e.g. fibroblasts/pericytes) in an extracellular matrix (ECM) precursor solution (e.g., fibrin, collagen) before seeding into the central channel of a microfluidic device, where flanking side channels are filled with culture media. Over the course of 5-7 days, endothelial cells form network structures that morph into open-lumen microvessels with surrounding pericyte-like cells, capturing architectural features of native blood vessels^1,5,7,8^. Flow is commonly induced by gravity, using rocking, or reservoirs with different media heights. Commercially available chips made from thermoplastics, which resist absorption of lipophilic media components, have popularized this approach^4,9^.

**Figure 1.**
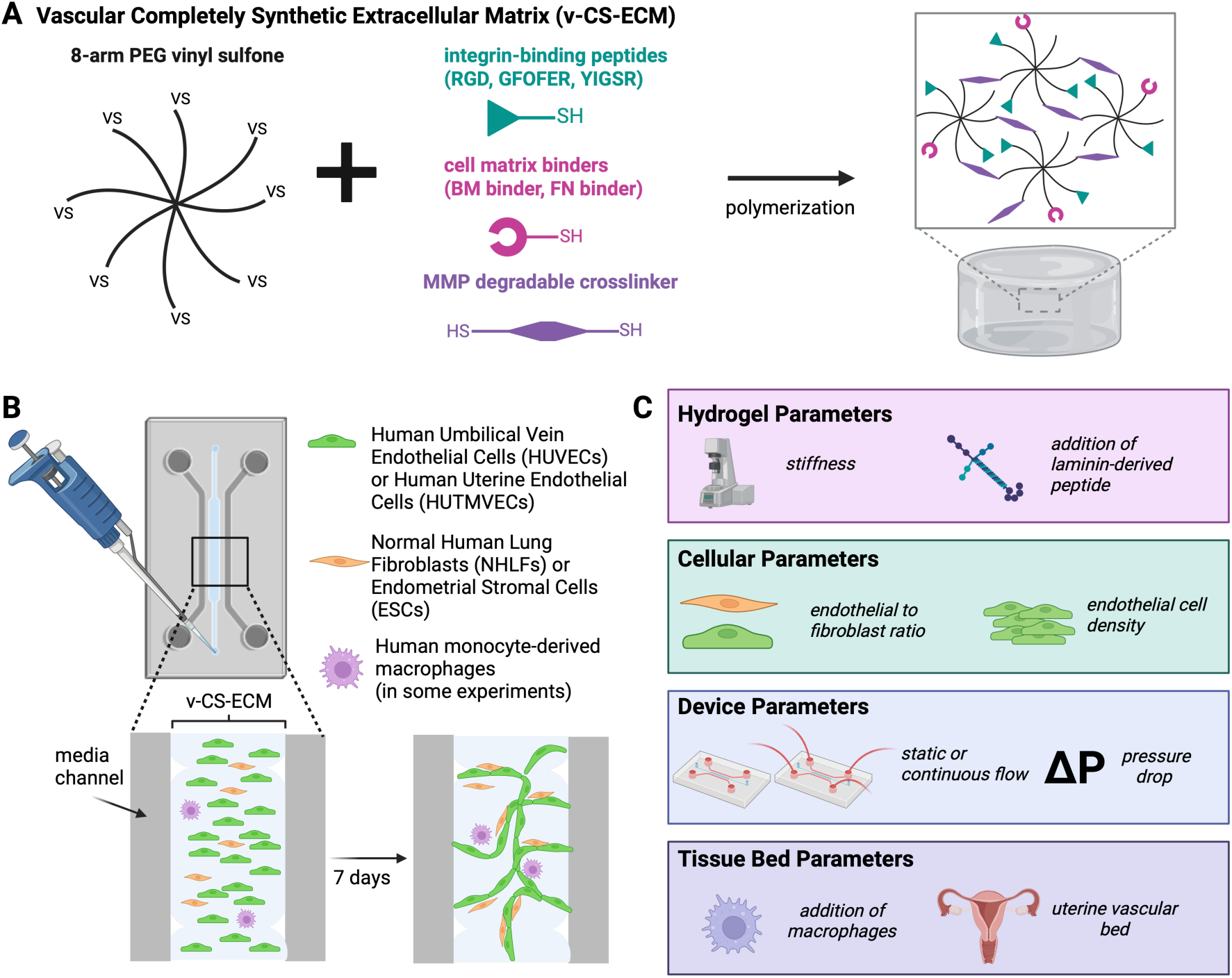
Overview of Synthetic Hydrogel Optimization for Perfusable Microvasculature. A) The v-CS-ECM comprises an 8-arm PEG-VS functionalized with integrin-binding and cell matrix-binding peptides and crosslinked with an MMP-degradable peptide for cell-mediated remodeling. B) The v-CS-ECM is mixed with endothelial cells and supporting cells prior to injection in a microfluidic device and vascular self-assembly over the course of a week. C) Hydrogel, cellular, device, and tissue bed parameters were varied to determine the impact on network formation in the v-CS-ECM.

Microvascular morphogenesis protocols rely heavily on fibrin, which serves as a provisional wound-healing ECM *in vivo.* As the expectations for reproducibility and biological performance of MPS continue to rise, the shortcomings of fibrin -- including lot-to-lot variability, undesirable biochemical cues, limited modularity, and poorly-controlled degradation^10–12^ – are becoming increasingly apparent. These shortcomings are accentuated in applications that incorporate additional cell types to create functional tissue mimics, as fibrin may impair or otherwise not support biological functions. For example, human mucosal epithelia, such as gut and endometrium, are expanded and cultured as organoids in basement membrane-mimicking Matrigel, which shares many of fibrin’s performance limitations but offers a complex mix of biological cues that support epithelia^13–15^. While a combination of fibrin and Matrigel enables a limited vascularization of gut organoids^16^, a more reproducible and biologically versatile ECM for creating complex microvascularized tissues is arguably desirable for MPS translation into the drug development pipeline. However, no completely synthetic extracellular matrix (CS-ECM) for vessel formation by morphogenesis *in vitro* has yet been reported^3–5^.

Building on design principles we developed for creating a peptide-modified polyethylene glycol (PEG) CS-ECM that supports endothelial network formation^11,17^ and endometrial epithelial-stromal co-cultures^18,19^, we recently described a CS-ECM formulation that is not only suitable for expansion and culture of primary patient-derived human epithelial gut, endometrial, and pancreatic organoids^20–22^, but also robustly supports co-culture of these organoids with stromal and immune cells^21,22^ in static (droplet) culture, where the organoids may stay entirely encapsulated or expand in size to emerge from the gel, forming a mucosal barrier along the surface of the CS-ECM^21^. The CS-ECM incorporates both synthetic peptides that bind to adhesion receptors known to be expressed by the target cells, as well as peptides that bind to native ECM produced by each cell type in the culture so that the cell-specific ECM is sequestered in the cell microenvironment^19,22^. This CS-ECM is crosslinked with matrix metalloproteinase (MMP)-susceptible peptides, enabling local cell-mediated remodeling. As with other synthetic or semi-synthetic ECM approaches^23^, the modularity of this organoid-supporting CS-ECM enables its properties to be tailored across a wide spectrum of biophysical, degradation, and compositional property ranges to mimic features of disease or control tissues and to support a wide range of co-cultured cell types^20–22^.

Here, inspired by the pressing need for humanized models of endometriosis lesions, we report design and function of the first CS-ECM that supports morphogenesis of perfusable microvascular networks in microfluidic devices, and demonstrate this formulation is compatible with expansion and culture of endometrial epithelia and stroma, and with creation of lesion-like structures surrounded by microvasculature. Endometriosis is a debilitating gynecological disorder that afflicts ∼10% of women and girls^24,25^.

Endometriosis lesions are defined as ectopic growth of endometrial-like tissue comprising endometrial epithelia and CD10+ stroma, typically comprising a polarized epithelial monolayer surrounding a fluid-filled lumen, supported by vasculature, nerves, and immune cell infiltrate^26^. As endothelial-fibroblast-epithelial interactions are crucially intertwined in determining how endometrial cells respond to changing hormone levels throughout the menstrual cycle^27–29^, an *in vitro* model system that captures these microvascular interactions, together with a perfusable vasculature to enable immune cell trafficking in long-term (weeks) culture, is highly desirable for analyzing patient-specific disease features and testing therapeutics.

The design approach and testing paradigm for this new “vascularizable” CS-ECM (termed here “v-CS-ECM”) (**Fig. 1A**) is guided by a semi-empirical screen of hydrogel and cellular parameters starting with those governing interconnected endothelial network formation in static microfluidic culture (**Fig. 1B-C**). Following analysis of outcomes in static culture, we introduce continuous fluid flow, systematically varying parameters to achieve lumenized endothelial network structures, termed vessels. Finally, we demonstrate this hydrogel is able support microvessel co-culture with organoids and macrophages, opening potential for complex disease modeling.

## 2. Results

### 2.1. Semi-Empirical Screen of Cell and Hydrogel Parameters Impacting Vascular Network Formation in Static Culture

#### 2.1.1 Introduction to Screen and Description of Microfluidic Device

We used a semi-empirical screen of various hydrogel, cellular, device and tissue bed parameters to accomplish microvessel morphogenesis in a CS-ECM hydrogel (**Fig. 1C**) to create v-CS-ECM, starting with morphogenesis of network structures in static culture, with a standard cell source, then progressing to perfusable vessels in cultures exposed to continuous flow and a tissue-specific cell source. While microvascular phenotypes are well-known to be tissue-specific^30^, practical issues of accessibility (including of cells already labelled with fluorescent reporters) and wide-spread use in other angiogenesis assays, contributed to the adoption of human umbilical vein endothelial cells (HUVECs) combined with normal human lung fibroblasts (NHLFs) as a workhorse model system for self-assembled microvessel formation^1,3,5,31^. As our ultimate goal is to develop protocols that can readily be adapted to a range of microvascular cell sources, we implemented a strategy of using HUVEC/NHLF in initial studies in screens as a benchmark against established work in the field, then extending the protocols to microvasculature cells relevant for endometriosis lesions.

Based on previous protocols described for vascular morphogenesis in microfluidic devices^3,32,33^, we designed and fabricated a custom device that could accommodate the anticipated relatively large size of a growing endometriosis lesion, with a vascular bed 1.8mm in width and 0.45mm in height (see detailed dimensions in Fig. S1; for reference, commercial AIM Biotech chips are 1.3mm wide and 0.25mm height). Four microposts on each side of the central gel channel pin the liquid hydrogel precursor solution via surface tension during loading. The chips were fabricated from PDMS, with appropriate pre-saturation with hormones where needed (Fig. S13).

#### 2.1.2 Hydrogel Composition and Biophysical Parameters

We reasoned that strategic modification of our CS-ECM*^21^* to include endothelial-targeted peptides, along with possible modulation of biophysical properties, would lead to a v-CS-ECM supportive of perfusable vasculature while retaining features supporting epithelial, stromal, and immune cell co-culture. In a previous semi-empirical screen of factors influencing endothelial cell network formation in PEG-based CS-ECM, we found that endothelial cells favor soft matrices*^11^*. However, an additional constraint is gel swelling post-crosslinking, as formulations that support network formation in free-swelling gels with large swelling coefficients may fail to support network formation in microfluidic devices, where swelling is constrained*^11^*. Therefore, combining these considerations, we aimed for a softer gel with a smaller swelling factor compared to the canonical regime used for organoids*^21^* by decreasing the weight percent of PEG and increasing the crosslinking ratio, respectively. Additionally, we have previously determined threshold concentrations of adhesion and matrix-targeting peptides required to support robust organoid and stromal co-culture (1.5mM PHSRN-K-RGD, 1.5mM GFOGER, 0.5mM BM Binder, 0.5mM FN Binder)*^20,21^*, hence kept these constant while adding a peptide targeted to endothelial interactions.

Laminin, an ECM protein, is an abundant component of the basement membrane in blood vessels*^34,35^*. Endothelial cells cultured in collagen hydrogels supplemented with laminin exhibit greater end-to-end endothelial network formation and increased VEGF uptake, compared to endothelial cells cultured in collagen alone*^36^*. Short adhesive peptides derived from laminin, including IKVAV from the laminin α1 chain and YIGSR from the laminin α1 chain are attractive candidates for creating v-CS-ECM *^37–39^*. YIGSR has been shown to regulate endothelial cell morphology in synthetic matrices, and when combined with RGD peptides, increases endothelial cell migration rates and endothelial tube length *in vitro^35,37,38,40^*. Given that we wanted to maintain the adhesion and matrix-targeting peptides developed for organoid culture, including RGD, we chose to incorporate YIGSR instead of IKVAV, as the combination of YIGSR and RGD in PEG hydrogels has been shown to form more stable endothelial tubule networks with increased tubule length and diameter compared to PEG hydrogels functionalized with a combination of IKVAV and RGD*^37^*.

As a starting point, we compared a softer hydrogel formulation (3.0wt% PEG, 0.52 thiol-ene crosslinking ratio) to a previous formulation used for organoids (3.6 wt% PEG, 0.43 thiol-ene crosslinking ratio). These initial experiments to screen hydrogel composition and biophysical parameters were performed with a total cell concentration of 12 x 10^6^/mL (7:1 HUVEC:NHLF), based on the upper end of the range of optimal results reported in fibrin. We used network area fraction, determined by a MATAB-based network quantification pipeline*^41^* of day 5 images (Methods) as a metric that captures significant differences between formulations, as network metrics can be reasonable predictors of later success in forming lumenized, perfusable vessels.

We found that this new formulation slightly increased the network area fraction in HUVEC/NHLF seeded into devices (from 0.30 to 0.34; **Fig. 2A**). This formulation demonstrated a storage modulus of ∼180Pa and a swelling factor of 1.5 (approximately 25% softer, and with 30% less swelling compared to the previous organoid formulation*^21^*, Fig. S2). Additionally, when we added YIGSR (0.6mM), while maintaining the concentrations of the four other peptides in the canonical CS-ECM constant, we found a significant increase in area fraction of endothelial cells compared to the previous organoid formulation*^21^*(from 0.30 to 0.38, p<0.05; **Fig. 2A**). Importantly, the addition of YIGSR did not change the hydrogel swelling or bulk stiffness (Fig. S2), indicating this increase in endothelial area fraction can likely be attributed to the peptide addition and not other hydrogel parameters.

**Figure 2:**
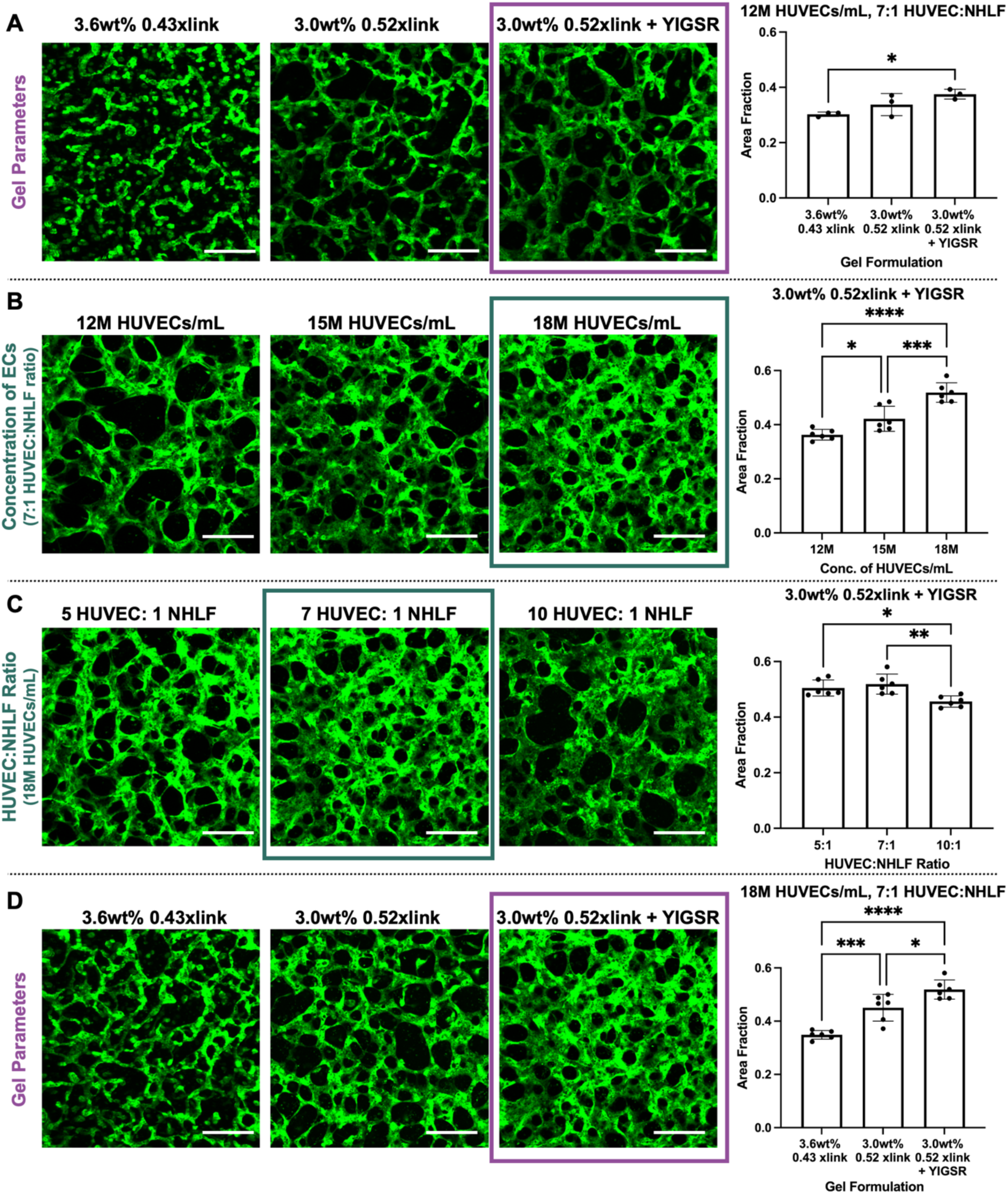
Hydrogel and Cell Parameter Static Endothelial Network Screen. A) Using the upper end of concentrations reported for fibrin (12M HUVECs/mL) a softer hydrogel (3.0wt% PEG 0.52xlink) compared to the previously published organoid formulation (3.6wt% PEG 0.43xlink) results in greater endothelial network area fractions which is further enhanced with the addition of YIGSR. B) Using the v-CS-ECM (3.0wt% PEG 0.52xlink + YIGSR), 18M HUVECs/mL resulted in significantly greater network area coverage compared to 12M and 15M HUVECs/mL. C) With 18M HUVECs/mL a 7:1 ratio of HUVECs to lung fibroblasts results in the greatest network area coverage. D) Using 18M HUVECs/mL and 7:1 HUVEC:NHLF ratio the difference in network formation between gel formulations is more pronounced, with the v-CS-ECM creating a significantly greater network area fraction compared to the same stiffness hydrogel without YIGSR (3.0wt% 0.52xlink) and the published organoid formulation (3.6wt% PEG 0.43xlink). Scale Bar: 200μm. Each data point represents a device. Max intensity projections from Day 5 shown. Data represented as mean±standard deviation. Statistics: ANOVA with Tukey post-hoc tests. *p<0.05, **p<0.01, ***p<0.001, ****p<0.0001.

#### 2.1.3. Cell Parameters

For the HUVEC/NHLF combination in fibrin, both formation of connected microvessel networks and ultimately perfusable microvessels depends on both the absolute and relative endothelial cell and stromal cell concentrations^3^. Networks formed in the optimal v-CS-ECM formulation described above were not perfusable, with a standard dextran perfusion assay initiated via gravity driven flow after 7 days of static culture. We therefore speculated that (i) the diameter of vessels formed was too low (<25μm) to support perfusion and immune cell circulation and (ii) morphogenesis in v-CS-ECM over the same time scale may require a greater number of endothelial cells compared to the standard concentration range (5-12M HUVECs/mL) needed for fibrin*^3,42^*.

To that end, we performed a semi-empirical parameter screen with 12-18M HUVECs/mL and ratios of 5:1, 7:1, and 10:1 endothelial to fibroblast cells. We determined that 18M HUVECs/mL with a 7:1 HUVEC:NHLF ratio resulted in the greatest area fraction in static culture conditions and used that combination for the remainder of studies involving HUVEC-NHLF cocultures (**Fig. 2B,C**). Notably, when using this optimized concentration and ratio, the v-CS-ECM formulation significantly increased the area fraction of endothelial cells compared to both the softer CS-ECM (0.45 to 0.52, p<0.05) and organoid formulation*^21^* (0.35 to 0.52, p<0.0001) (**Fig. 2D**, Fig. S3). This condition formed interconnected networks with an area fraction of ∼0.5 and a mean network diameter approximately 35μm; however, despite significant network interconnectivity which is an important first step towards creating vessel-like structures, these networks still did not have open-lumen structures necessary to enable perfusion. We therefore reasoned that, as *in vivo*, fluid flow factors may play a crucial role in development of fully lumenized, perfusable vessels.

### 2.2 Introducing Flow and a Pressure Drop Results in Perfusable Vasculature

Vessels *in vivo* are constantly exposed to hemodynamic forces, which influence their remodeling and functions^10,10,43^. Many investigators use gravity-driven pressure gradients generated by a height difference in media reservoirs feeding the side channels^44^. Here, to avoid flow transients associated with gradual equilibration of reservoir heights, we adapted the static culture device to configurations incorporating recirculating peristaltic pump-driven flow through the side channels, using the return channel loop to generate defined weak or strong pressure drops across the tissue compartment, as either a simple loop or a serpentine return (**Fig. 3A**, Fig. S4). In these flow setups, the flow rate was controlled at 1μL/s, imposing a shear stress on the hydrogel walls of approximately 0.07 dyne/cm^2^ (Fig. S5). Importantly, all three device designs (static, simple-loop, serpentine-loop) have the same tissue chamber geometry and dimensions (**Fig 3A**, detailed dimensions in Fig. S1), isolating continuous flow and pressure difference as the only changing variables.

**Figure 3:**
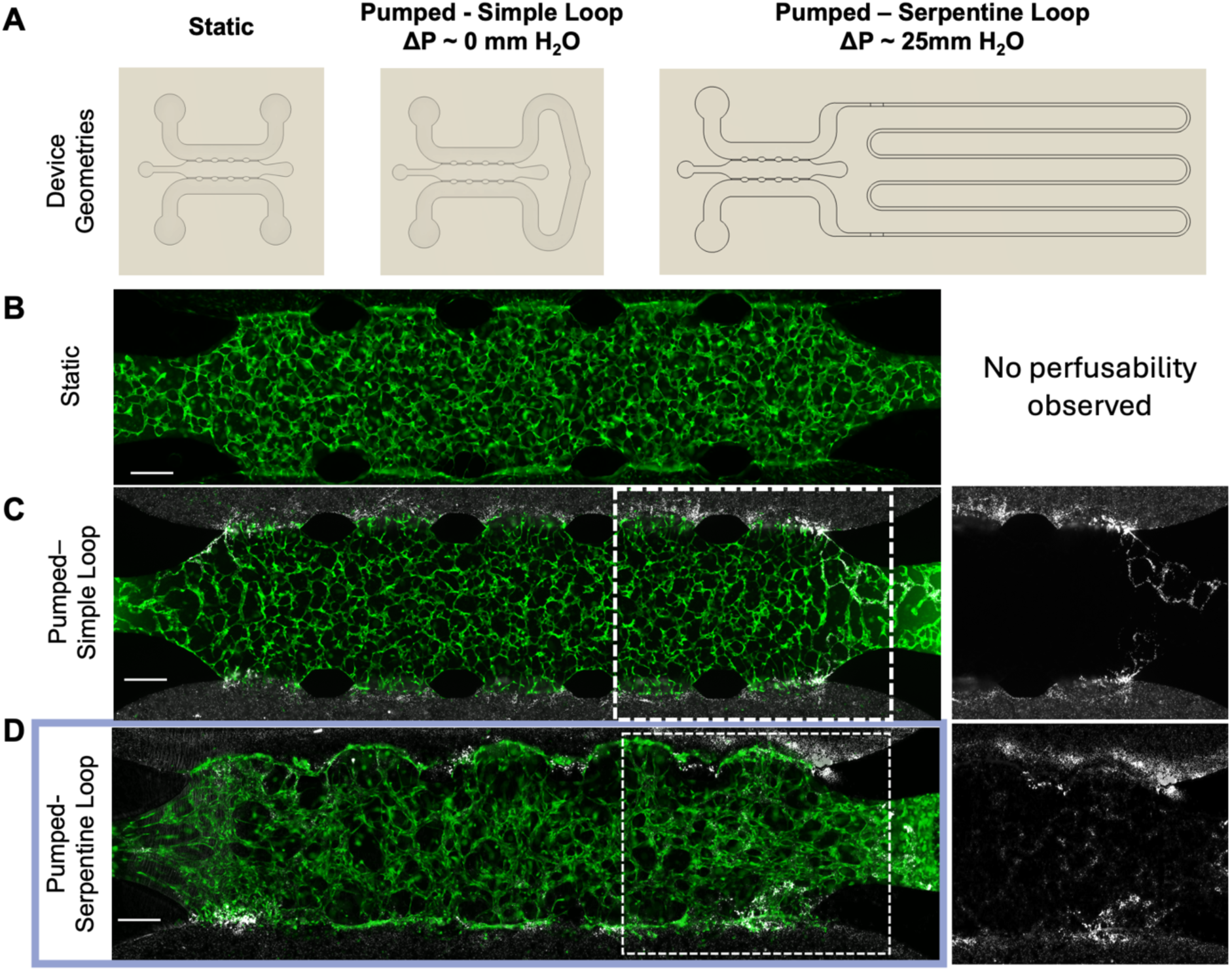
Transition from static to continuous flow culture results in perfusable microvasculature. A) As we transitioned from static culture conditions to continuous flow driven by peristaltic pumps, the device geometry was adapted to have a simple looped flow or serpentine looped flow that had a measured pressure difference of 25mm H_2_O across the channel. B) Full device view of networks cultured in static conditions show an interconnected network without perfusability observed. C) Full device view of networks cultured in a looped device without a pressure differential demonstrated perfusability at the end of the channel evidenced by beads trafficking through networks (zoomed in bead only view in the white boxed area). D) Full device view of networks cultured in a serpentine device with a pressure differential demonstrated perfusability across the channel evidenced by beads trafficking through networks (zoomed in bead only view in the white boxed area). Scale Bar: 500μm.

In the first configuration, we used a simple loop generating a very small (0.2 mm H_2_O) ΔP (**Fig. 3A**, Fig. S5), as this would emphasize the imposition of shear rather than transmural flow. After culturing for 7 days, vessels near the inlet and outlet gel-injection ports displayed open lumens and were perfusable as assessed via the flow of microbeads (**Fig. 3C**), though networks near the center of the device did not become perfusable.

We hypothesized that a more significant pressure drop across the channel may increase the fraction of perfusable vessels, as the biophysical forces imposed by pressure drops influence endothelial cell physiology^45,46^ and are commonly implemented in vascularized MPS^44^. To increase the ΔP in the context of a constant imposed recirculating flow, we implemented a serpentine looped channel design (**Fig. 3A**). We measured the trans-tissue-channel ΔP as approximately 25mm H_2_O (Fig. S4) close to the value of 32mm H_2_O predicted by a COMSOL model (Fig. S5). Notably, when we cultured microvessel networks using the serpentine device exposed to flow throughout the entire duration of the experiment, we observed perfusable vessels, evidenced by the flow of fluorescent microbeads through open-lumen structures across the channel (**Fig. 3D**, **Fig. 4A**). The fraction of vessels was approximately 45% (**Fig. 3D**, Fig. S6), with most perfusable vessels in the top quarter and bottom quarter of the device. Additionally, these vessels could be maintained in culture for at least 14 days under continuous flow (Fig. S7). As an attempt to decrease the high concentration of endothelial cells required, we tested 15M HUVECs/mL (7:1 HUVEC:NHLF ratio) and observed perfusable vessels in the bottom quarter of the device (Fig. S8), indicating a partial success and there are likely other parameters that could be optimized to decrease the endothelial cell density. While further iteration on parameters such as side channel geometry (influences shear stress on surface endothelia) and post placement (influences tissue biomechanics via its effects on gel compression and on durotaxis) is needed to reach 100% perfusable vessels across the entire channel, the current findings are a significant advance in the development of a completely synthetic hydrogel for microvascular morphogenesis. We did not pursue extensive further iterations at this point as applications, such as the endometriosis lesion model we aspire to, involves additional cell types that will likely influence remodeling and vessel maturation. We therefore proceeded to investigate both underlying mechanistic hypotheses for differences in performance in these configurations, as well new phenomenology that might arise from inclusion of additional cell types of interest in applications.

**Figure 4:**
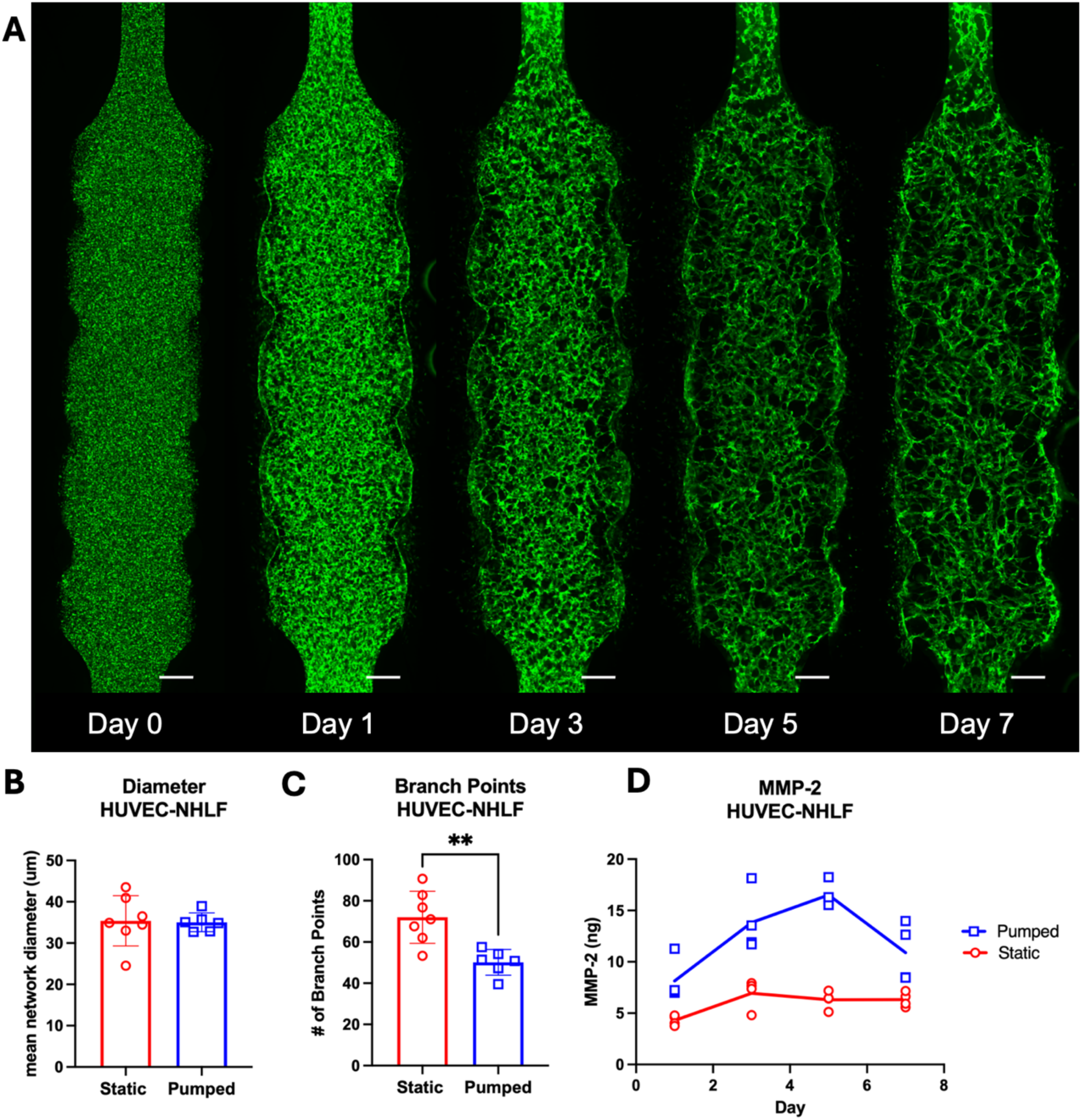
Vessel Formation in continuous flow conditions. A) Self-assembly of HUVEC-NHLF networks over 1-week results in perfusable vessels in a device with continuous flow and a pressure differential. B) Diameter does not significantly change between static and pumped conditions. C) Network complexity assessed by the number of branch points is significantly reduced in flow conditions. D) Pumped cultures have significantly greater MMP-2 secreted in spent medium compared to static culture conditions, corresponding with greater perfusable network formation. Each data point represents a device. Scale Bar: 500μm. Data represented as mean±standard deviation. **p<0.01.

### 2.3 ECM Remodeling Contributes to Perfusable Vessel Formation

MMP-mediated ECM remodeling is one potential flow-induced mechanism contributing to open lumen formation in our synthetic hydrogel between static and continuous flow conditions. MMP-2 has been shown to facilitate vessel formation in fibrin hydrogels^47^, and the MMP-sensitive crosslinker in v-CS-ECM is susceptible to degradation via MMP-2. We therefore measured MMP-2 concentrations via ELISA on media samples collected after 24 hr of circulation, at the time of daily media changes, throughout the time course of the morphogenesis process (days 1, 3, 5, and 7). Strikingly, there were significantly higher MMP-2 levels detected via ELISA in the spent medium from networks cultured with continuous flow compared to those networks cultured in static conditions (**Fig. 4D**). The MMP-2 levels were strikingly higher at days 3 and 5 in the pumped condition compared to static and started to decrease at day 7. We have observed the open lumen structures emerging between day 5 and day 7 (**Fig. 4A**), and hypothesize this enhanced remodeling contributes to the endothelial network to vessel progression. Therefore, this could be a useful marker for evaluating future hydrogel formulations for microvessel formation, as we found this elevated MMP-2 corresponded with open lumens that were not present in static conditions. Unexpectedly, we did not observe a discernible difference in network diameter between static and serpentine-looped pumped devices (35μm for both static and pumped conditions, **Fig. 4B**), demonstrating an increased vessel diameter did not always enable perfusable vessels. We note that lack of perfusion does not necessarily mean that vessels lack lumens; lumenized vessels may fail to open up to the side channels. The pumped devices fostered simpler network structures evidenced by significantly fewer branch points (**Fig. 4C**, static = 72, pumped = 50, p<0.01), which has been similarly observed for HUVEC-NHLF cultures in fibrin hydrogels^43,48^.

### 2.4 Successful Incorporation of Macrophages into Endothelial Networks

While the bedrock of microvascular engineering focuses on co-culture of endothelial cells with a fibroblast population, macrophages are key regulators of vessel sprouting, serve as sources of angiogenic growth factors, and are observed to align with nascent vasculature similar to support cells^49^. Macrophages are phenotypically plastic innate immune cells ubiquitous to tissue microenvironments, where they perform tissue homeostatic functions including lipid and iron processing, along with wound-healing and immune defense against pathogens^50,51^. The homeostatic tissue-resident macrophage population is augmented when needed by trafficking of circulating CD14+ monocytes into the tissue, driven by chemokine cues such as CCL2^50,51^, contributing further to inflammatory cascades that influence vessel and tissue function^52,53^. Macrophages are implicated as primary drivers of endometriosis lesion pathophysiology, and are viewed as a target of therapeutic intervention^25,54–56^. We therefore investigated protocols for incorporating macrophages.

To generate macrophages, CD14+ human monocytes were isolated from healthy donors and differentiated over 6 days using 50 ng/mL macrophage-colony stimulating factor. We implemented the static culture screen (described above, and Methods), starting with keeping a constant concentration of HUVECs at 18M/mL, while varying the concentration of macrophages in the supporting cell population (macrophages, NHLFs) from 0-100%, keeping the ratio of endothelial: supporting cells at 7:1 and the total concentration of supporting cells at 2.57M/mL (**Fig. 5A**).

**Figure 5:**
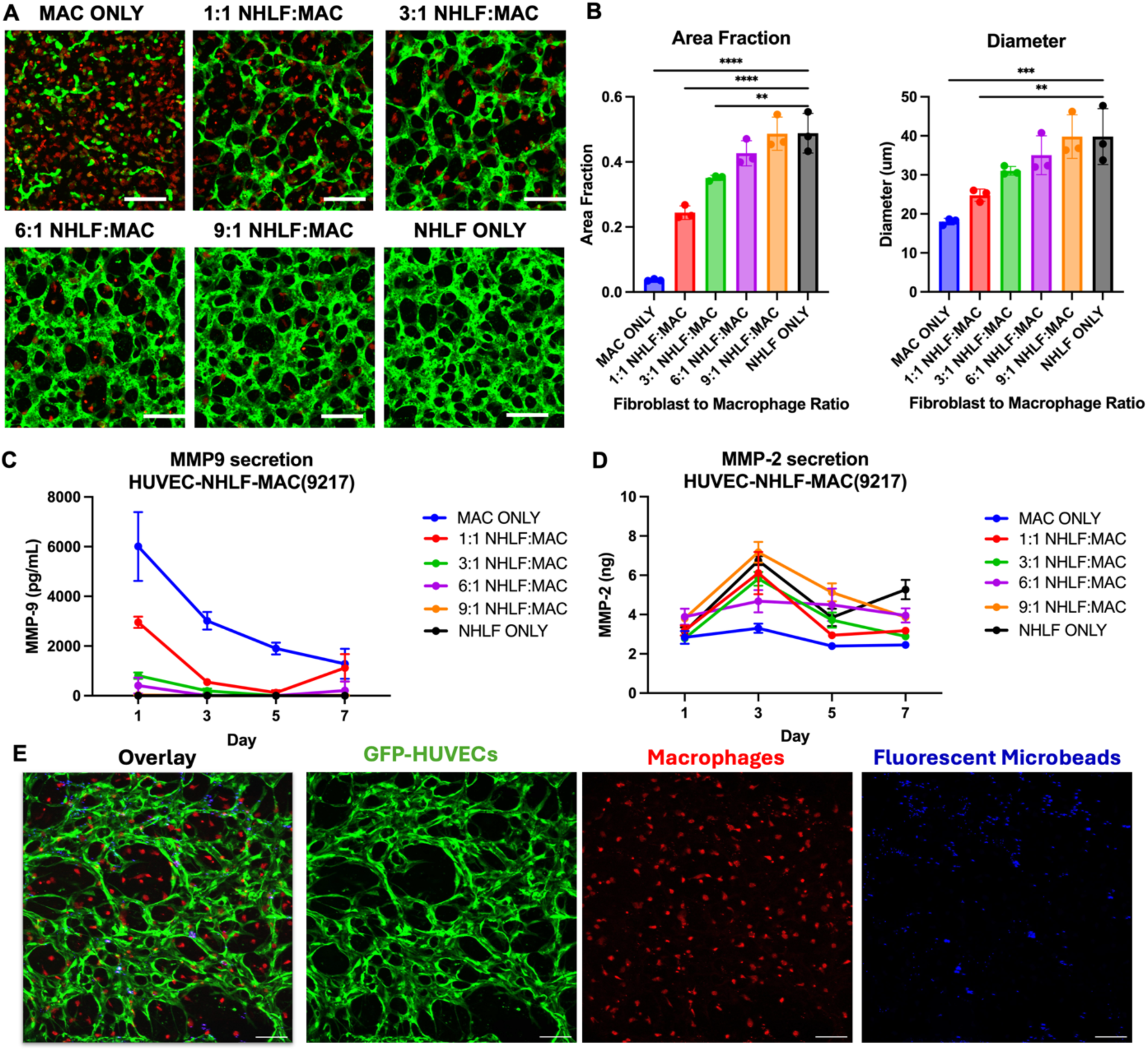
Incorporation of macrophages in HUVEC-NHLF networks. A) Day 5 confocal images of HUVEC-NHLF-macrophage networks in static culture with varying ratios of fibroblasts to macrophages (macrophages in red labeled with CellTrace). B) As the macrophage ratio increases the area fraction and diameter of the endothelial networks decreases. C) Macrophages are functional in culture and secrete MMP-9. D) MMP-2 secretion over 7-day static culture in HUVEC-NHLF-macrophage culture. E) Transitioning the 6:1 NHLF to macrophage ratio to pumped culture, the HUVEC-NHLF-macrophage networks are perfusable as evidenced by bead trafficking through the endothelial networks (green). Scale Bar: 200μm. Statistics: ANOVA, Tukey post-hoc test. *p<0.05, **p<0.01, ***p<0.001, ****p<0.0001. Data represented as mean ± standard deviation. N=3 devices.

As the percentage of macrophages increased from 0-100% the endothelial network diameter, area fraction, and length density decreased (**Fig. 5B**). Cultures entirely lacking NHLF (100% macrophage in supporting cell population) exhibited scattered network fragments but did not generate fully formed networks. The 9:1 and 6:1 fibroblast: macrophage ratios supported endothelial network diameter and area fraction values comparable to controls with no macrophages. In day 5 imaging (**Fig. 5A**), macrophages appeared to have heterogeneous spreading morphologies with clear projections, an observation corroborated by Moore et. al who previously showed that macrophage morphology is highly spread in endothelial-macrophage co-cultures in a PEG-based hydrogel^57^.

To confirm the functionality of macrophages in our system, we analyzed media supernatants for MMP-9, a major MMP produced by human macrophages that is known to have an influence on angiogenesis^46,58,59^. The amount of MMP-9 secreted corresponded with the number of macrophages in the culture, and was highest at day 1, with no detectable MMP-9 in the NHLF only condition (**Fig. 5C**). The endothelial network morphology and MMP-9 trends were repeated with a second macrophage donor, giving us confidence macrophages maintain functionality in the v-CS-ECM (Fig. S9). We similarly analyzed MMP-2 and did not see a clear trend with the presence of macrophages, except the macrophage only condition had the lowest total secreted enzyme (**Fig. 5D**).

Once we established that macrophages could be incorporated into microvascular networks, we confirmed the ability to form perfusable vessels with the 6:1 macrophage ratio. Similar to the HUVEC-NHLF only networks, perfusable networks formed in the serpentine loop devices under continuous flow, as evidenced by beads trafficking through the networks (**Fig. 5E**) and likewise observed the significant increase in MMP-2 secretion when transitioning from static to pumped culture (Fig. S10). We observe close proximity between the macrophages and vessels, likely indicating the macrophages are serving in a supporting cell capacity as previously reported in the literature^49,60^. Together, this model provides a launch point to study the influence of innate immune cell behavior on vascular networks in a disease-specific context.

### 2.5 Uterine Microvascular Endothelial Cells form Perfusable Vessels in v-CS-ECM

While HUVECS and NHLFs are a useful system to benchmark vascular morphogenesis, vascular phenotypes vary significantly between tissues^30,45,61,62^, prompting us to evaluate how well protocols for the benchmark cell types can be extended to microvasculature relevant for a disease-specific model. Adenomyosis is a sibling disease of endometriosis, characterized by endometriosis-like lesions in the myometrium of the uterus. Noting that the fibroblasts in lesions have characteristics of endometrial stromal cells^63^, we purchased commercial uterine microvessel endothelial cells, and combined them with primary patient-derived endometrial stromal cells (ESCs) as a model for microvasculature in the vicinity of an adenomyosis lesion.

In similar fashion to our initial cell parameter screen with HUVEC and NHLF cell concentrations and ratios (**Fig. 2B,C**), we performed a parameter screen with varying concentrations of endothelial and stromal cells. We again found 18M HUTMVECs/mL and a ratio of 7:1 to result in the greatest network area fraction (0.41) and diameter (31μm) (**Fig. 6A,B**). When exposing these cultures to continuous flow, vessel structures were perfusable as evidenced by beads trafficking through the open lumens across the device (**Fig. 6C**). Strikingly, the average vessel diameter was significantly increased (36μm, **Fig. 6D**) upon the transition from static culture to continuous flow conditions, as has been observed previously for generic vasculature formed with HUVECS/NHLFs in addition to brain-specific microvasculature in fibrin hydrogels^10,43,48,64^. This observation is in contrast to the HUVEC/NHLF case, where the transition from network features observed in static to flow were limited to simpler network structures, without a significant change in vessel diameter (**Fig. 4B-C**).

**Figure 6:**
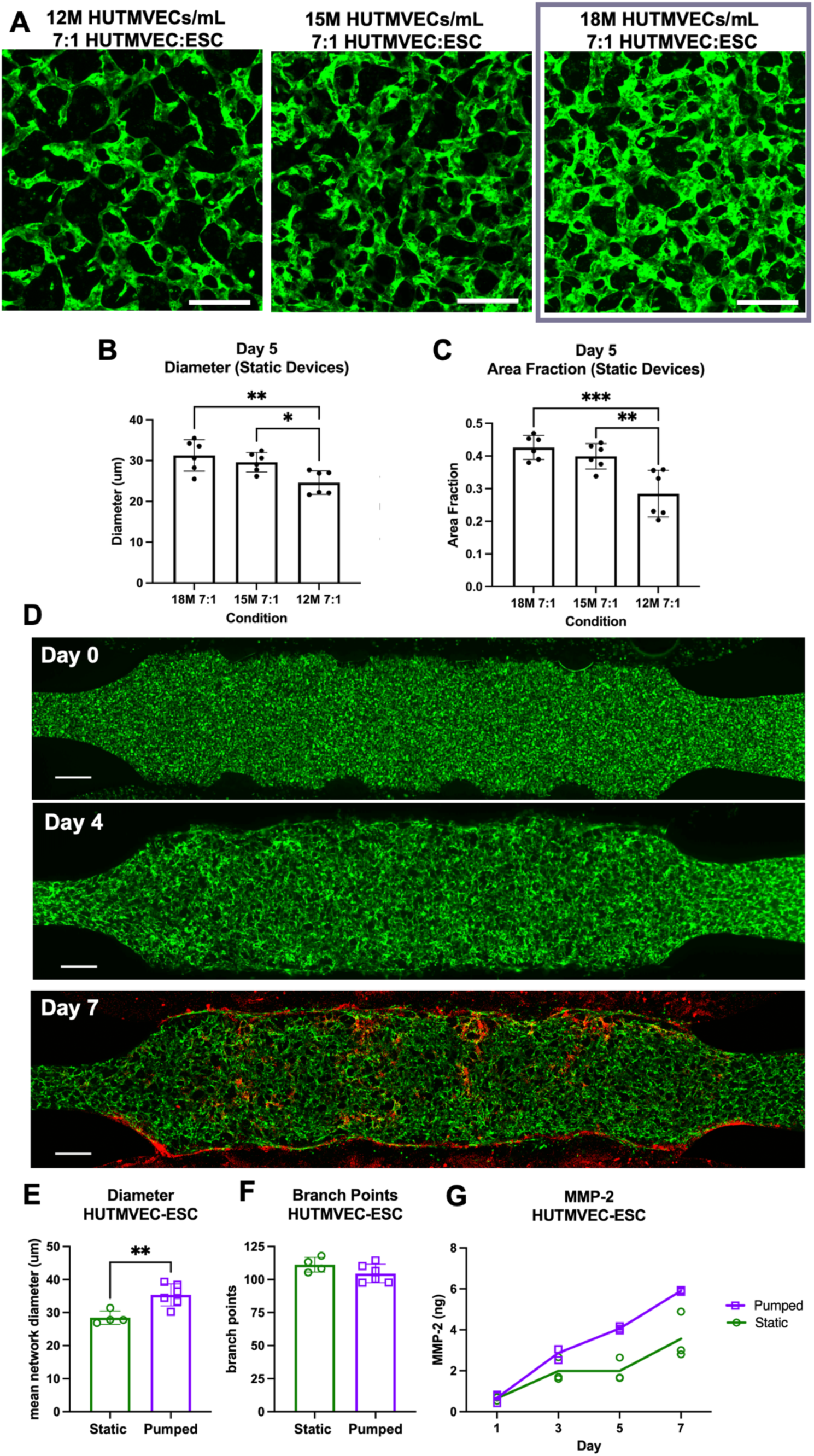
Uterine Endothelial Networks. A) Day 5 images of GFP-labeled uterine endothelial cell (HUTMVEC) – endometrial stromal cell (ESC) networks in static conditions. B) The same concentration of endothelial cells and ratio of endothelial cells to fibroblasts (18M 7:1) resulted in the greatest network area coverage and network diameter. C) HUTMVEC-ESC networks were exposed to continuous flow in a serpentine-loop device and self-assembled into perfusable vessels over a course of 7 days as seen by red beads trafficking through the networks (labeled with GFP) at Day 7. D) In pumped conditions the vessel diameter was significantly greater compared to static, E) there was not a significant difference in number of branch points representing network complexity, and F) more MMP-2 was detected over time in spent culture media. Each data point represents a device, N=3-6 devices. Scale Bar: 200μm. *p<0.05, **p<0.01, ***p<0.001, ****p<0.0001. Data represented as mean ± standard deviation.

Mirroring previous observations there is a significant increase in MMP-2 detected in the spent media from flow conditions compared to static culture (**Fig. 6E**), however the relative levels of MMP-2 are much lower for the uterine microvessel cell source than from HUVEC/NHLF. Additionally, there is a continuous increase in MMP-2 throughout the course of a weeklong culture, potentially pointing towards the slightly slower network formation observed via daily imaging with these cells compared to HUVEC-NHLF coculture. These results were repeated with a second ESC donor with no significant changes in outcomes (Fig. S11).

### 2.6 Co-culture of Endometrial Epithelial Organoids with Uterine Microvascular Networks

Endometriosis lesions are defined as ectopic growth of endometrial-like tissue, typically comprising a polarized epithelial monolayer surrounding a fluid-filled lumen with surrounding CD10+ stroma, supported by vasculature and immune cells^63,65^ (**Fig. 7A**). After demonstrating that the v-CS-ECM can support self-assembly of perfusable vessels formed with uterine-derived endothelial cells and ESCs (**Fig. 6**), we investigated creation of an endometriosis lesion model by adding patient-derived endometrial epithelial organoids to the cell population at the time of culture initiation.

**Figure 7:**
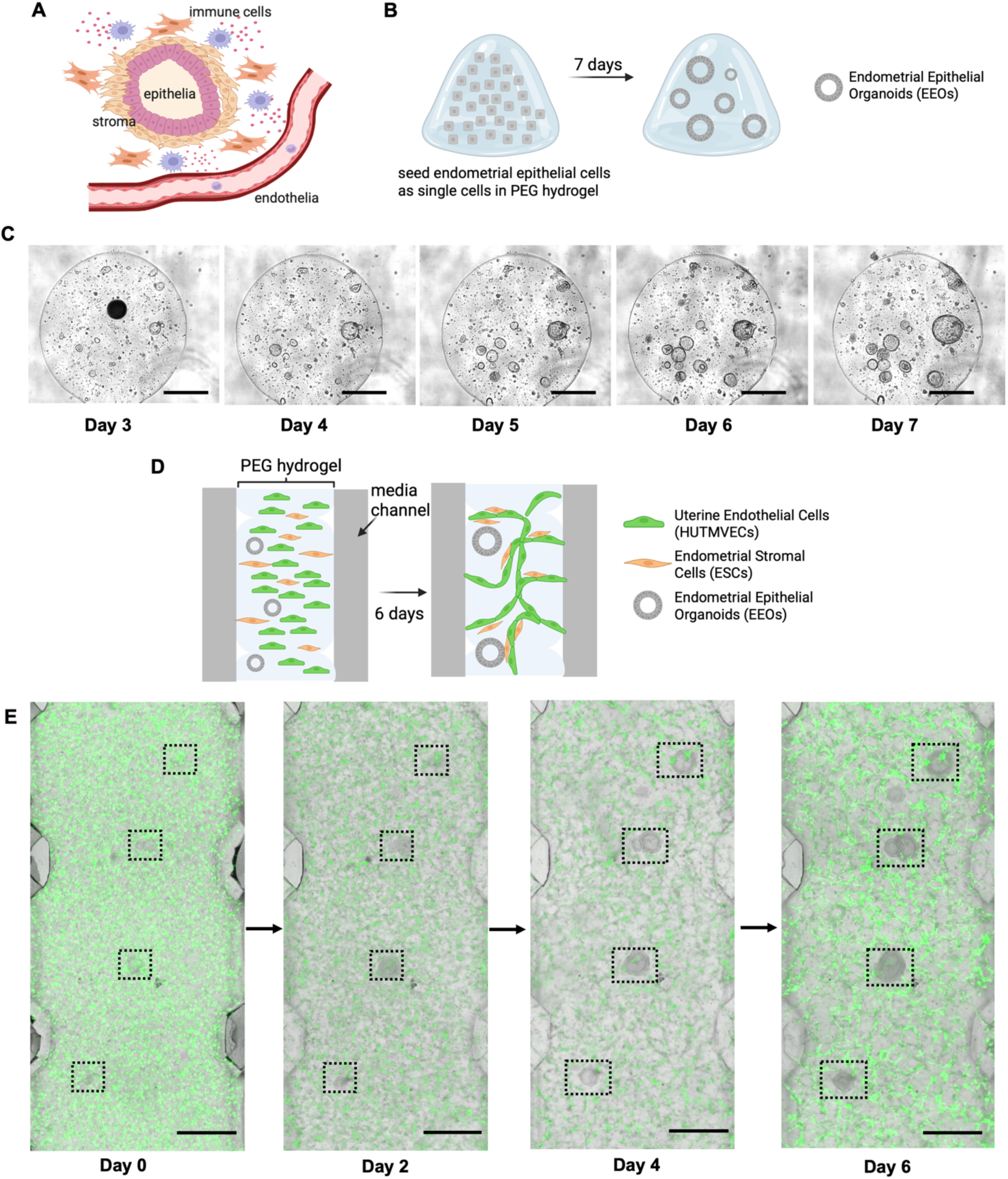
Incorporation of Organoids in Uterine Microvascular Networks. A) Endometriosis lesions establish complex microenvironments consisting of epithelial, stromal, endothelial, and immune cells. B) Droplet hydrogel cultures were used to assess organoid emergence from single cell. C) v-CS-ECM supports organoid emergence from single cell over the course of a week. D) EEOs, HUTMVECs, and ESCs were combined in a hydrogel precursor prior to injection in a microfluidic device and cultured for 6 days. E) V-CS-ECM supports EEO growth within HUTMVEC-ESC networks. Scale Bar: 500μm.

Prior to combining vasculature and organoids, we first assessed primary endometrial epithelial organoid (EEO) emergence from single cells encapsulated in v-CS-ECM using hydrogel droplet culture (**Fig.7B**). Similar to the canonical CS-ECM we previously described for EEO culture^21^, the v-CS-ECM supports robust EEO emergence from single cells (**Fig. 7C**). As a consideration before combining all cell types relevant for co-culture in the full lesion model, a suitable mixed media formulation was developed to support both the EEOs and vasculature. A 50-50 mixture of standard microvascular endothelial media was mixed with our organoid media^21^ excluding the TGFα inhibitor A8301 and Rock inhibitor Y-27632^21^. This media was shown to support network formation in static culture to a comparable extent to standard endothelial cell media (Fig. S12). Additionally, endometrial EEOs are typically cultured in an estradiol (E2) containing medium, which is rapidly absorbed into PDMS. To mitigate this, we developed a presoaking method to saturate the PDMS with estradiol and maintain the desired concentration over the duration of the experiment (Fig. S13). We note that for conditions simulating a full menstrual cycle, where hormone concentrations must be cycled, PDMS devices offer formidable challenges and use of relatively non-absorptive thermoplastics is desirable.

Having established that v-CS-ECM supports EEO expansion, patient-derived EEOs approximately 100μm in diameter were combined with patient-matched ESCs and commercially sourced uterine endothelial cells into the hydrogel precursor solution (**Fig. 7D**), then injected into a serpentine-looped microfluidic device connected to continuous flow. Over the course of approximately a week, the organoids grew significantly in size as the microvessels self-assembled similar to previous experiments (**Fig. 7E**, Supplemental Video 1). This demonstrates the v-CS-ECM and microfluidic device developed in this manuscript can support the coculture of epithelial, stromal, and endothelial cells. Notably, this system enables the formation of lesion-like structures and mirrors the structure of *in vivo* lesions where vessels do not penetrate the lumens of endometriosis lesions but instead surround the fluid-filled epithelial monolayer structures, modeled by EEOs in this system^63^. Therefore, our model is physiologically appropriate when it excludes vessels from within the center of the lesion and has them surrounding the lesion. While the results shown here are simply a proof-of-principle experiment showing this v-CS-ECM enables growth of lesion-like structures in the microfluidic devices, this sets a launching point for more developing a better scientific understanding of endometriosis and can be used in the future for testing novel drug compounds.

## 3. Discussion

There has been substantial literature focusing on the use of fibrin and collagen hydrogels to create self-assembled vasculature enabling foundational studies on vessel biology. Despite significant efforts towards creating self-assembled microvascular networks in synthetic hydrogels, it has been very difficult to achieve microvessels with open lumens which can serve as viable replacements for fibrin or collagen^5,11,66^. Some success has been achieved within a sprouting angiogenesis microfluidic model, with reports of a dextran-based hydrogel that can support open lumen formation after a 21-day window^66–68^. Additionally, hyaluronic acid-based hydrogels supporting endothelial tubules have been reported, with microscopic evidence of lumens and basement membrane structures, however these culture models were not reported in microfluidic devices^69,70^.

Establishment of a synthetic hydrogel to support perfusable vasculature will further increase the utility of vascularized MPSs in the regulatory space as synthetic PEG-based hydrogels are highly reproducible, modular, and can serve be tailored to specific cell or tissue applications^12^. Our synthetic hydrogel platform was designed specifically to enable *in vitro* modeling of the local pericellular microenvironments each cell type in a disease state experience, as cells can deposit their own ECM and remodel the matrix into a tissue-relevant ECM^19,21,22^.

In the body, vasculature is subject to fluid flow and mechanical stresses as new vessels are developing and throughout homeostasis^71,72^. The incorporation of flow to vascularized microphysiological systems further improves the physiological relevance of these model systems, and flow has been shown to play a critical role in aiding in vessel formation and maintenance *in vitro*^43,47^. Upon introduction of flow and a pressure drop within a normal physiologic range of capillaries and previously reported microfluidic models^44,73^, we were able to achieve vessels with open lumens in our v-CS-ECM in microfluidic devices. Of note, our estimated wall shear stress on the hydrogel is significantly lower than what capillaries experience *in vivo* (∼40 dyn/cm^2^)^74^ and future studies are needed to investigate how increasing the shear stress to a physiologically relevant range would influence vessel formation.

To investigate a potential mechanism to explain why vessels had open lumens when exposed to flow but remained networks in static culture, we examined mechanisms governing extracellular matrix remodeling. The MMP-cleavable crosslinker in our v-CS-ECM is susceptible to degradation by MMP-2 and MMP-9, both of which have known associations with vascular remodeling^46,47^. Throughout the experiments presented in this manuscript, networks that had been exposed to continuous flow had significantly increased MMP-2 levels in media supernatant collected daily compared to static conditions, and this was observed for both HUVEC-NHLF and HUTMVEC-ESC networks. This finding has also been observed in fibrin hydrogels, with a recent study showing that vessels were unable to form when exposed to an MMP-2 inhibitor. While higher concentrations in the pumped case compared to static may arise from enhanced convective transport of MMP-2, in fibrin, MMP-2 was upregulated upon exposure to interstitial flow and plays a significant role in vessel formation^47^. Future studies can address regulation of MMP-2 expression at the transcriptomic and tissue level. Additionally, when we added macrophages into the system we found they secreted MMP-9, which was not detected in HUVEC-NHLF networks without macrophages. This can further contribute to the degradation of our hydrogel, though high concentrations of macrophages limited network formation, and more work is needed to determine exact mechanisms governing perfusable vessel formation in our synthetic ECM.

An open question is the physical mechanism by which a pressure drop across the tissue compartment influences gene expression and function in v-CS-ECM compared to fibrin, as v-CS-ECM has a very tight pore structure with extremely low hydraulic conductivity, such that negligible fluid flows across these gels in the absence of cells, whereas fibrin has a more open pore structure and a measurable hydraulic conductivity that allows interstitial flow rates to be achieved at the pressure drops we use. A plausible hypothesis about fluid flow through the v-CS-ECM gels is that cell migration in the gels opens continuous conduits within the first few days. In addition to the obvious morphological changes of the endothelial cells pictured in Figs. 4, 6, and 7, fibroblasts that are not visible are also migrating. We have previously observed tracks of laminin left by migrating iPSC-derived endothelial cells in similar gels^11^, after the cells had moved on to elsewhere in the gel. Such ECM tracks suggest open channels were left behind by the cells after migration through the gel.

Another drawback of current vascularized MPSs is the primary reliance on HUVECs and supporting fibroblasts typically sourced from the lung^2^. While these cell types are easy to obtain and create perfusable microvascular networks, microvascular cells from adult organs display different structural attributes and molecular expression profiles^2,30^. As more vascularized MPS models have been developed, there are starting to be more tissue-specific endothelial cells in perfusable microvascular models, including mammary microvessels^45^, brain microvessels^3,75^, and dermal microvessels^3^.This is the first report to our knowledge of perfusable uterine endothelial cells self-assembled in a microfluidic device, with limited research utilizing uterine-specific endothelial cells^76–78^. We are actively using this uterine bed vascular model for endometriosis lesion modeling, with the first proof of principle experiments shown here (**Fig. 7**) and anticipate this v-CS-ECM will support other tissue-specific endothelial cells, opening avenues for tissue-specific vascularized models.

There are a few limitations of this model system, including that the networks must be exposed to flow to become perfusable, unlike fibrin hydrogels, which can limit the throughput of experiments. However, demonstration of partially perfusable self-assembled networks is a significant improvement compared to current synthetic hydrogels for vascularized applications and introduction of flow aids in physiological relevance. Additionally, while we briefly determined a biochemical mechanism contributing to open-lumen vessel formation in this paper, future studies are needed to better parse the effects of ECM composition, cellular composition, mechanical cues, device design, and flow profiles on vessel formation in the v-CS-ECM. Finally, for the purpose of modeling endometriosis and the complex hormonal dynamics exhibited throughout the menstrual cycle, we recognize the need to shift to thermoplastic devices. For the purposes of this work focusing on optimizing the v-CS-ECM for perfusable vessel formation, PDMS devices were used as they are amenable to quick fabrication and design changes. Active efforts are focused on creating thermoplastic alternatives with similar device geometries developed in this foundational work for downstream disease modeling and drug testing, and future work will focus on extended culture timelines mimicking the different hormonal regimes of the menstrual cycle.

Overall, this paper demonstrates a new tool that can be used in vascularized models with room for many future explorations including the combination of organoids and perfusable vasculature for complex MPS.

## 4. Conclusion

In this paper, we report development of the first synthetic ECM that supports perfusable self-assembled vascular networks in microfluidic devices. We first modified our existing CS-ECM formulation that has been shown to support organoids^20,21^, by decreasing the stiffness and adding YIGSR, a laminin-derived peptide. We empirically screened the number of endothelial cells and supporting fibroblasts to support a robust interconnected endothelial network in static culture conditions before introducing continuous fluid flow. With continuous flow and exposure to a continuous pressure difference across the tissue channel, we achieved vasculature with open lumens in a fully synthetic hydrogel. Additionally, we confirmed the ability of this hydrogel to support addition of macrophages to endothelial networks and support tissue-specific microvascular endothelial cells, setting a launch point for complex disease modeling. Finally, we demonstrate this matrix can support the coculture of vasculature and epithelial organoids using endometriosis lesion modeling as a preliminary validation. Overall, we anticipate this hydrogel should have far reaching impacts on the microfluidic community by creating a reproducible, fully defined matrix to support vascularized tissue models.

## 5. Experimental Section/Methods

### Preparation of Hydrogel Components

8-arm 20kDa poly(ethylene glycol) (PEG) macromers functionalized with vinyl sulfone (PEG-VS) were purchased from JenKem Technology (Beijing, China). All peptides were custom synthesized and purified (>95%) by CSBio (Milpitas, CA), CPC Scientific (Sunnyvale, CA), or GenScript (Piscataway, NJ). Peptides used in these studies include: a dithiol crosslinking peptide with an MMP-cleavable substrate (Ac)- GCRDLPRTGGPQGIWGQDRCG-(Amide) (**LW**); a fibronectin-derived peptide containing the canonical RGD motif as well as the PHSRN synergy site previously described^21^, (Ac)-PHSRN- GGG-K(GGGERCG-Ac) GGRGDSPY-(NH_2_) (**PHSRN-K-RGD**); a collagen-derived peptide, (Ac)-GGYGGGPGGPPGPPGPPGPPGPPGFOGERGPPGPPGPPGPPGPPGPC-(NH_2_) (**GFOGER**); a peptide with affinity for sequestering basement membrane proteins type IV collagen and laminin, (Ac)-GCREISAFLGIPFAEPPMGPRRFLPPEPKKP-(NH_2_) (**BM Binder**); a peptide with affinity for sequestering cell secreted fibronectin, (Ac)- KKGCREPLQPVYEYMVGV-(Amide) (**FN Binder**); and a peptide derived from binding motif of laminin, (Ac)-GCREYIGSR-(Amide) (**YIGSR**). PEG-VS was dissolved in sterile Ultrapure DNase/RNase-Free Distilled Water (Invitrogen) using a volumetric flask to create a 10% w/v solution in a biosafety cabinet and aliquoted and stored at -80°C until use. All peptides were reconstituted in sterile Ultrapure DNase/RNase-Free Distilled Water (Invitrogen) in a biosafety cabinet at a target concentration of 30mM for PHSRN-K-RGD, GFOGER, YIGSR and LW; 15mM for FN binder; 20mM for BM binder. To fully dissolve YIGSR a small amount of DMSO (<10% of total volume) was added. To fully dissolve FN binder the pH must be adjusted to a final pH of 5-7 using a concentrated NaOH solution. Peptide solutions were sterile-filtered, aliquoted, and stored at -80°C until use in the gel. All aliquots were used for less than 3 freeze-thaws. Peptide concentrations were measured using Ellman’s assay (Sigma 22582) for GFOGER, 205nm absorbance for PHSRN-K-RGD, LW, BM binder, and FN binder, and 280nm absorbance for YIGSR.

### Cell Culture

GFP-labeled Human umbilical vein endothelial cells (HUVECs) and GFP-labeled human uterine microvascular endothelial cells (HUTMVECs) were purchased from Angio-Proteomie (cAP-0001GFP and cAP-0023GFP). Normal Human Lung Fibroblasts (NHLFs) were purchased from Lonza (CC-2512). Endometrial stromal cells (ESCs) and endometrial epithelial organoids (EEOs) were from our endometrial tissue bank and isolated and expanded using previously published methods^21^ under IRB-P0012994. HUVECs were cultured in Vasculife VEGF Endothelial Medium (Lifeline Technologies LL-0003), HUTMVECs were cultured in Vasculife VEGF-Mv Endothelial Medium (Lifeline Technologies LL-005), and fibroblasts were cultured in Fibrolife S2 Fibroblast Medium (Lifeline Technologies LL-0011). ESCs were cultured in DMEM/F12 with 5% charcoal-stripped fetal bovine serum (FBS, Biotechne #S11650), 1% Penstrep (Gibco 15140122), and 1nM Estradiol (Sigma-Aldrich E8875). EEOs were cultured in Matrigel Growth Factor Reduced, Phenol Red-free (Corning 356231) prior to using in our synthetic hydrogel in a media previously described^21^. All cell types were used between passages 5-9.

### Monocyte isolation

Human monocytes were isolated from healthy donors, either isolated from buffy coats obtained from Massachusetts General Hospital or from peripheral blood Leukopaks (StemCell Technologies 200-0092). CD14^+^ monocytes were isolated from buffy coats as previously described^79,80^. First, peripheral blood mononuclear cells (PBMCs) were isolated via density gradient centrifugation with Lymphoprep (STEMCELL 18061). PBMCs were washed with MACs buffer: PBS supplemented with 2% charcoal-stripped FBS (Biotechne #S11650) and 2 mM EDTA (Corning 46-034-CI), and isolated using a EasySep Human CD14 Positive Selection Kit II (STEMCELL 19359) according to the supplier’s instructions. Leukopaks were processed similarly without the initial density gradient step. Once isolated, CD14+ cells were cryopreserved at 10 million cells/vial in freezing media (90% charcoal-stripped FBS with 10% DMSO).

### Macrophage differentiation

Cryopreserved CD14+ monocytes were thawed into RPMI 1640 media (Gibco 32404014) supplemented with 10% charcoal-stripped FBS, 1% Pen-Strep (ThermoFisher 15140148), GlutaMAX (ThermoFisher 35050061), then seeded in dishes at a density of 1 million cells per mL in 10 mL of complete media with 50 ng/mL human M-CSF (BioLegend 574804). Cells were incubated at 37°C/5% CO_2_ for a total of 6 days, with a media change on day 3. To harvest cells for downstream experiments, cells were washed with PBS and incubated in MACs buffer at 4°C for 20 min followed by gentle washing with a p1000 before collection and centrifugation at 300 G for 5 min. Macrophages were stained by incubating with CellTrace™ Far Red Cell Proliferation Kit (Invitrogen C34572, 1:1000) in a 50:50 solution of MACs buffer and PBS for 20 min at 37°C and washed with 25X volume of MACs buffer before cell counting for downstream experiments.

### Hydrogel Creation

Synthetic hydrogels were assembled using PEG weight% and crosslinking ratios denoted throughout the text. In all studies, PHSRN-K-RGD and GFOGER were used at 1.5mM and BM binder and FN binder used at 0.5mM. YIGSR was added to the vascular formulation at a concentration of 0.6mM. PEG-VS concentrations was12mM for the 3.0wt% condition and 14.4mM for the 3.6wt% condition. To create the hydrogel, first PEG-VS macromers were functionalized with the monothiol pendant peptides (PHSRN-K-RGD, GFOGER, BM binder, FN binder, YIGSR) in a buffered solution, with 10% of the volume 10x PBS + 1mM HEPES, pH 8.2. Briefly all components were added at appropriate concentrations in sterile conditions, vortexed thoroughly, and allowed to react via a Michael-type addition for 30 minutes at room temperature. If including cells (all studies except rheology in Fig. S2), the desired cells were lifted and pelleted using standard protocols during the 30-minute functionalization step. Cells were mixed at the desired concentration for a certain volume of gel (ex. 18M HUVECs/mL, 7:1 ratio – for a 40μL gel this is 720,000 HUVECs and 102,857 NHLFs in a pellet) and pelleted using centrifugation (300gx5min) in a microcentrifuge tube. Carefully, all media was removed from the cell pellet after centrifugation, and the pre-functionalized PEG solution was added to the cell pellet at an appropriate volume and mixed via gentle pipetting at least 15 times. Next, quickly the LW crosslinker was added to create a desired thiol-ene crosslinking ratio of 0.43 or 0.52 to the PEG-cell mixture. Again, the gel is thoroughly mixed using pipetting at least 15 times prior to loading the pre-gel solution into the device using a 10μL pipet. Gel batches were typically prepared in 40μL volumes to load 3-4 devices per batch. After loading, devices are placed at 37°C in a humidified incubator for 30 minutes to allow gelation to occur via a Michael-type addition. Media was added to each device after the 30-minute gelation incubation. For experiments including EEOs, the EEOs were grown in Matrigel to an average diameter of approximately 100-150μm before harvesting in Cell Recovery Solution (Corning 354253) to dissolve the Matrigel. Intact organoids were added at a concentration of 3 EEOs/μL of gel to the endothelial-stromal cell pellet before mixing with the functionalized PEG solution. *Rheology:* To measure the bulk mechanical properties of the hydrogel, 50ul of the pre-gel mixture was pipetted into 8mm silicone molds (0.5mm thick) which were stuck to slides that had been coated with SigmaCote (Millipore Sigma SL2) and allowed to gel at 37°C. After gelation, hydrogel discs were removed from the silicone molds/ glass slides using a plastic spatula and weighed using a balance to get the pre-swelling mass. The hydrogel disc was then placed in a 12-well plate with 2mL of PBS. The plate was incubated for 24 hours in a 37°C incubator to allow equilibrium swelling to occur prior to rheological testing. After swelling, the hydrogel discs were weighed on a balance to get the post-swelling mass to calculate the mass swelling factor. An Anton Paar rheometer (model MCR302) was used to measure rheological properties. The hydrogel discs were sandwiched between an 8mm sandblasted parallel plate and sandblasted stage set to 37°C. The shear storage modulus (G’) was measured by performing a time sweep for 180s at 1% strain and 1 rad/s angular frequency.

### Device Fabrication

Polydimethylsiloxane (PDMS, Sylgard 184 Silicone Elastomer Kit, Dow) devices were fabricated using a 10:1 ratio of silicone elastomer base to curing agent. Briefly the PDMS mixture is poured in a silanized stiff PDMS negative mold (5:1 ratio of silicone elastomer base to curing agent) and allowed to cure for at least 2 hours at 65°C. The PDMS negative molds are created from a silanized positive milled acrylic mold (CAD file available upon request). All devices have a central 1.8mm wide channel flanked by two media channels (Supplementary Figure 1). The gel-/media-channel interface features a 0.1mm phase guide and 8 posts (4 per side), which assist with proper hydrogel loading. Serpentine devices also feature a narrow, serpentine channel (0.3mm channel width) connecting both media channels. This narrow channel design creates a pressure drop across the gel channel of approximately 32mm H_2_O at a flow rate of 1 µL/s, calculated with COMSOL as described below.

1.2 mm holes were punched using biopsy punches at both ends of the gel channel in all devices. In static devices, four 6mm holes were punched at the ends of both media channels. Once bonded these large punches act as media reservoirs, allowing the device to hold 300µL of media per device. In serpentine devices, two 1.2mm holes were punched into the non-serpentine ends of the media channels. Once bonded sterile syringe tips are then inserted into these holes connecting the device to an external peristaltic pump and media reservoir.

Before bonding, all PDMS devices were washed sequentially with methanol, isopropanol, distilled water, and methanol, dried with an air hose, then autoclaved. Devices were bonded with the microfluidic channel side facing downwards, leaving only punched areas exposed. Static devices were bonded to sterile 6-well glass bottom plates (Cellvis P06-1.5H-N), while serpentine devices were bonded to sterile standard glass microscope slides. Every device was stored in an oven at 65°C for at least one day prior to use.

### Pumped Device Setup

All pump circuit components were sterilized according to the methods outlined in Supplementary Figure 4C. These components were then assembled inside a biosafety cabinet (Supplementary Figure 4D), with the media reservoir intentionally left disconnected. Pump tubing was then fed into an Instech P625 peristaltic pump (Instech) and held within the pump by a 3D printed pump cap. 0.5mL of media was then circulated through the pump circuit, allowing excess media to drain into a waste collection tube. After priming, the inlet and outlet connectors were attached to the media reservoir by the top port and side port respectively. An Eppendorf® protein LoBind tube (Millipore Sigma) with 1mL of media was also secured into the media reservoir.

To connect the serpentine devices, one tube was detached from each pump circuit’s syringe tip and affixed to a new, sterile syringe tip. These two syringe tips were then inserted into the pre-punched holes at the top of each device media channel. For the lifetime of each experiment, devices were stored inside PREempt-sterilized 3D printed imaging bases (as shown in the bottom image). Caps from three 15mL conical tubes were secured into the base and filled with PBS to keep the base chamber humidified, then a sterile universal 96-well plate lid was placed on top of the base to seal the chamber. The full system was then transferred to a standard cell culture incubator, and the pumps were subsequently powered to deliver a constant volumetric flow rate of 1 µL/s.

The imaging base was designed to fit inside a Keyence BZ-X710 imaging microscope (Keyence). Pump caps were required to keep the pump tubing from falling out of the pump during extended use of the peristaltic pumps. Both are made from polylactic acid. CAD files for both parts are available upon request.

### COMSOL Modeling of Pressure and Fluid Shear Stress on Hydrogel

The pressure difference and shear stress on the hydrogel was numerically simulated using COMSOL Multiphysics version 6.3. Briefly, a steady, laminar, single phase flow assumption was used based on the Reynolds number (∼1.5) in the media channels. The tissue chamber was not modeled as the flow through the hydrogel was assumed to be negligible, compared to the flow through the side channels. The shear stress at the interface between the hydrogel and media was calculated to be ∼0.07 dyne/cm^2^. This was calculated by multiplying the shear rate reported by the simulation with the dynamic viscosity of the fluid. The pressure difference between the two side channels was estimated to be ∼32 mmH_2_O, created by the hydraulic resistance to flow in the long serpentine connecting segment between the side channels. The fluid properties used in the simulation were taken from Poon^81^, for media containing 5% FBS (density 1002 kg/m^3^, dynamic viscosity 0.862 mPa.s).

### Pressure Drop Experimental Measurements

Vertical column pressure drop measurements (Supplementary Fig. 4A) were taken to verify that the serpentine narrow channel creates a pressure gradient across the hydrogel. For testing, pump circuits were set up identically to a normal experiment and connected to serpentine devices loaded with a cell-free PEG gel. Tubing in the pump circuit was cut immediately before the device inlet and after the device outlet to insert 1/16” tee union hose barb fittings (T-connector, Cole Parmer). Excess 0.031” I.D. Tygon tubing was attached to the third barb of the T-connector and was loosely taped to the wall so that the tubing created parallel vertical columns. Media was circulated through the device for 10 minutes before measuring the media height difference between the inlet and outlet vertical columns.

### Device Maintenance

Devices were imaged daily at 4x using a Keyence BZ-X710 microscope in the green channel. Media was changed daily, with 1mL of media exchanged on the pumped devices and 300μl of media for static devices (150μl per side). Media was collected daily in Nunc™ 96-well U-bottom plates (Thermo Scientific 268200) and centrifuged at 1000g x 5 min and then transferred to a new plate for storage at -80°C until downstream ELISA analysis.

### Media composition for experiments

In HUVEC-NHLF experiments (Fig. 2-4), devices were cultured with Vasculife VEGF Endothelial Medium (Lifeline Technologies LL-0003). In experiments containing macrophages (Fig. 5), the Hydrocortisone Hemisuccinate concentration in Vasculife was reduced to 500nM to reduce the effect of cortisol on macrophage activity. In HUTMVEC-ESC experiments without organoids (Fig. 6), devices were cultured in Vasculife VEGF-Mv Endothelial Medium (Lifeline Technologies LL-005). In HUTMVEC-ESC experiments with organoids (Fig. 7) a 50-50 mixture of VL VEGF-Mv Endothelial Medium (kit FBS was replaced with charcoal-stripped FBS) and Endometrial Epithelial Organoid medium^21^ without A8301 and Y-27632 with 10nM E2 was used for the duration of the experiment. In all experiments, media was changed and collected daily.

### ELISAs

Human MMP-2 (R&D Systems DY902) and Human MMP-9 DuoSet (R&D Systems DY911) ELISAs were performed on spent medium. Manufacturer protocols were adapted to be performed in a 384-well plate to minimize the sample volume needed. Standard curves were fit to a 4-parameter logistic curve using MATLAB or GraphPad Prism. Technical duplicates were performed for all samples and standard curves were performed in at least triplicate. MMP-2 values were normalized to total ng of MMP-2 to account for medium volume differences in static and pumped culture settings.

### EEO Emergence studies

EEOs were thawed into Matrigel and expanded as previously reported. Once the EEOs were lumenized with an average diameter greater than 250μm, the Matrigel droplets were harvested using cold Cell Recovery Solution (Corning 354253). Briefly, Matrigel droplets were collected with wide-bore pipette tips and mixed with Cell Recovery Solution and placed on ice for approximately 20-30 minutes until the Matrigel was fully dissolved. After dissolving, the solution was centrifuged at 300g x 5 min to get a cell pellet. Organoids were manually broken into single cells using previously described protocols^21^ in a TrypLE (ThermoFisher 12604021) solution containing DNAase (Sigma Aldrich D4527-40KU) with manual pipetting. Single cells were counted and resuspended in a functionalized PEG solution (all components except crosslinker) at a density of 1000 cells/μL of gel. Next, the LW crosslinker was added at the desired concentration and mixed thoroughly before pipetting 3μL droplets in a non-TC treated 96 well plate. The hydrogel droplets were placed in a 37°C incubator for 30 minutes to allow gelation to occur before adding 150μL of complete EEO medium (including ROCKi). Imaging was performed every day using a Molecular Devices ImageXpress for the course of a week to visualize EEO emergence. Media changes were performed every other day.

### E2 Presoaking

PDMS rapidly absorbs hormones, therefore, to mitigate this issue we developed a method to pre-saturate the devices with estradiol (E2) prior to using them in experiments. To do this, devices were fully submerged in a 10nM E2 solution in PBS for 24 hours and then the solution was aspirated, and devices were dried in a 37°C vacuum oven overnight. Controls for this experiment included devices that were not presoaked and devices that were presoaked and did not get media with E2. After presoaking devices (in a solution of 10nM E2 or PBS without E2) and drying, blank gel without cells was loaded into 3 devices per condition (9 devices total). After gelation, media containing 10nM E2 or without E2 was added and changed daily for a duration of 6 days. Media was collected each day and stored at -80°C for downstream analysis. After the experiment, media from each condition was analyzed using an estradiol ELISA kit (DRG International EIA2693). The concentration of estradiol in the spent medium was normalized to the 10nM E2 media to calculate the fraction of E2 that was taken up by PDMS. *Network Morphology Analysis:* Devices were imaged at either Day 5 or 7 on a Zeiss LSM 880 confocal microscope with a 10x objective using a 488 laser to image the GFP-labeled endothelial cells. Z-stacks are taken at a least 3 distinct sites per device and export a maximum intensity projection (MIP) for analysis. Image files are processed using REAVER^41^, a MATLAB based analysis, to quantify mean network diameter, total network length, network length density, and area fraction.

### Perfusability Quantification

Perfusability was assessed by the flow of fluorescent microbeads (Invitrogen FluoSpheres Polystyrene Microspheres, 1.0μm) which were added to the media reservoir and circulated for at least an hour on the final day of the experiment, with experiments ending between day 7 and day 10. Since the beads do not fully fill the vessels and flow across the vessels, perfusability was quantified per region of the device. If at least half of the vessels in each region of the device contained beads that section of the device was considered perfusable. Quantification was performed by dividing the device per post region and the 2 edges (Supplementary Figure 7B), with each post region representing 16.6% (1/6) of the total area and the edges together representing 16.6% (1/6) of the device.

### Quantification and Statistical Analysis

Data are expressed as mean± standard deviation, with individual data replicates shown. Each point represents a single device. Graphs were created and statistical analysis was performed using GraphPad Prism 10. Specific sample sizes are noted on each figure legend. Statistical significance is defined by *p<0.05, **p<0.01, ***p<0.001.

## Acknowledgements

This work was funded by National Institutes of Health U01EB029132, National Institutes of Health R01HD114214, a sponsored research agreement with NovoNordisk, Manton Foundation, Fairbairn DAF, and John and Karine Begg Fund. Schematics created with BioRender.com. The authors would like to thank Dr. Samantha Holt, Ellen Kan, and Erin Tevonian for technical expertise and support throughout this project.

## Author Contributions

L.P and L.G conceived the study. L.G.G. supervised the study. L.P designed experiments. L.P., and R.O. conducted experiments. L.B. helped design and conduct all experiments with macrophages. A.J. assisted with maintenance on experiments and performed ELISAs. P.R., M.J., and D.T. performed device design and fabrication. L.P. wrote the manuscript. All authors reviewed and provided feedback on the manuscript.

## Conflict of Interest

L.P. and L.G. have filed a patent application (U.S. Application No. 63/892,868) on the vascular synthetic hydrogel.

## Notes

### Competing Interest Statement

The authors have declared no competing interest.

