## Supplemental Figures for "Development of a Synthetic Hydrogel to Foster Microvascularization of an Endometriosis Microphysiological System"

Supplementary Figures:

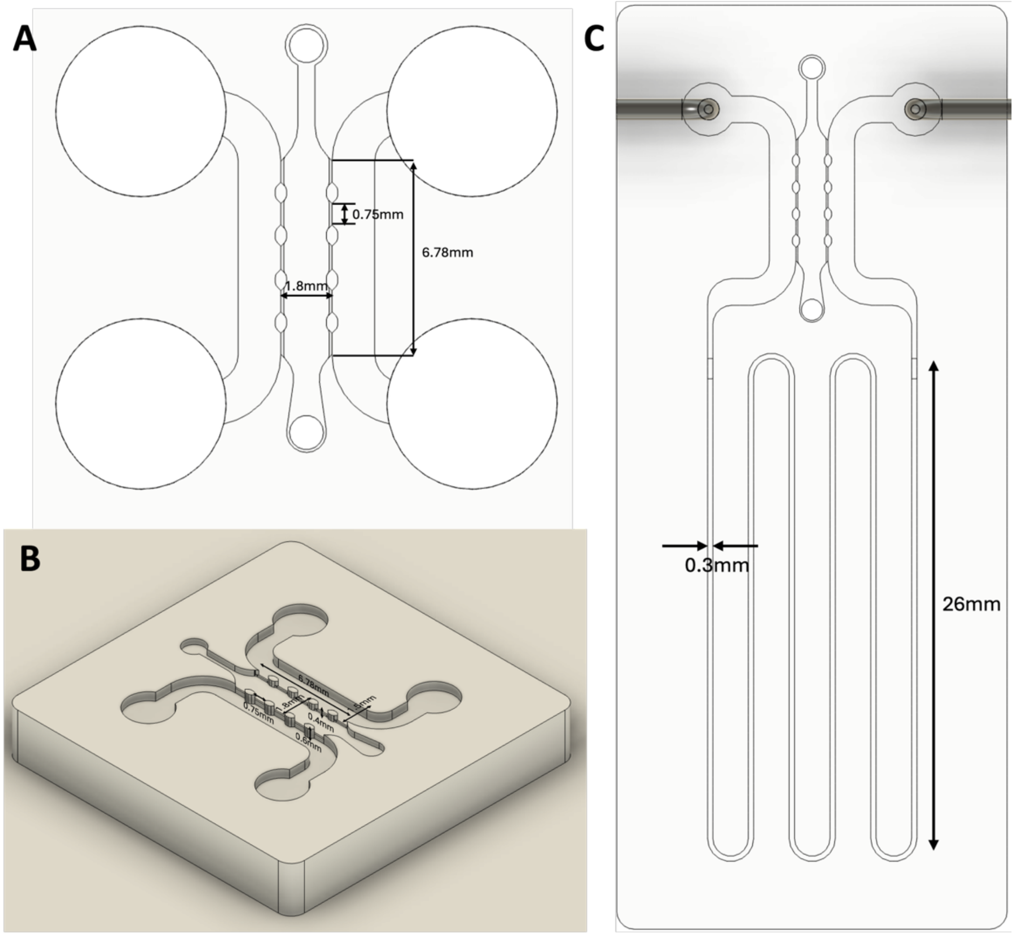


**Supplemental Figure 1: CAD snapshots of custom microfluidic devices.** A-B) Dimensions of the static device used in this study. C) Dimensions of the serpentine channel added to the device to enable flow studies with a continuous pressure drop. The tissue channel geometry is the same as described in A-B.

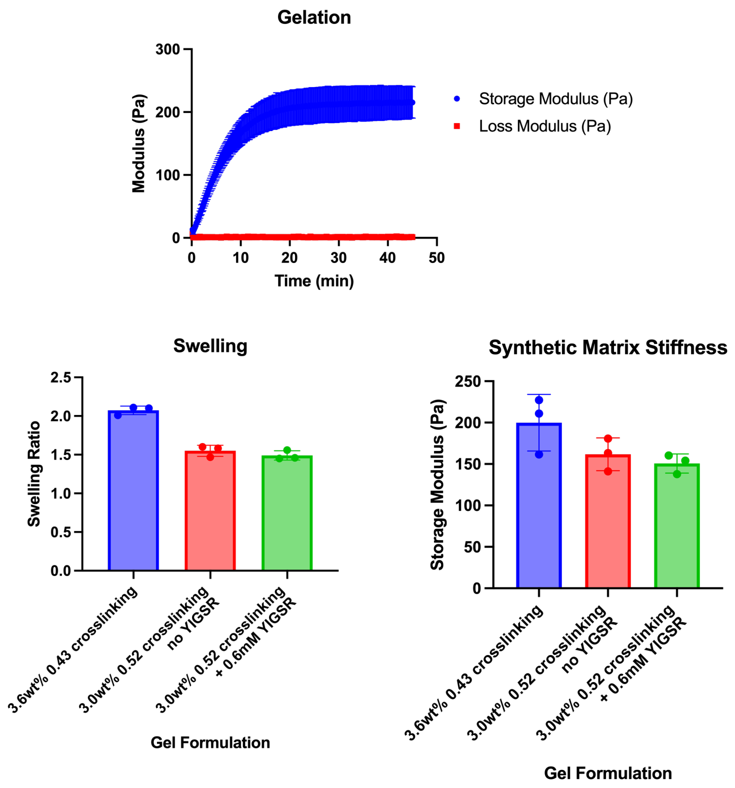


**Supplemental Figure 2:** A) In situ gelation curve for the 3.0wt% 0.52 crosslinking hydrogel with YIGSR shows gelation is fully complete after 30 minutes at 37°C. B) Mass swelling ratio for the three hydrogel formulations tested in this paper. The 3.0wt% 52% crosslinking hydrogel formulations swelled ~25% less compared to the original organoid formulation published in Gnecco et al[21]. C) Bulk stiffness measurements using rheology to compare the three gel formulations. The vascular formulation was ~150Pa storage modulus and 25% softer.

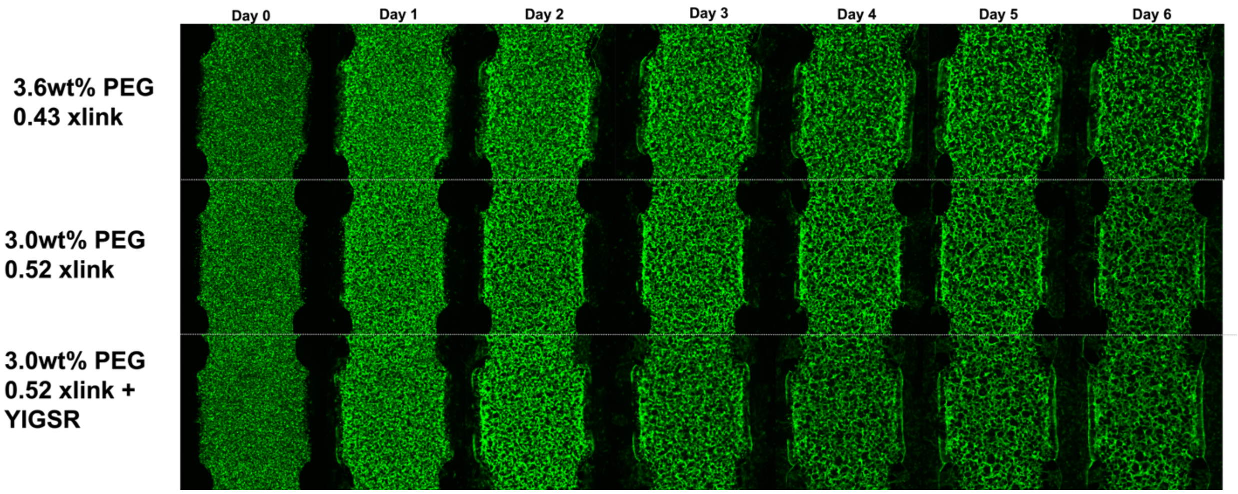


**Supplemental Figure 3:** Daily device imaging to monitor network progression in each of the different gel formulations used in this manuscript cultured in static conditions. Concentration of HUVECs: 18M/mL and 7:1 HUVEC:NHLF ratio.

A

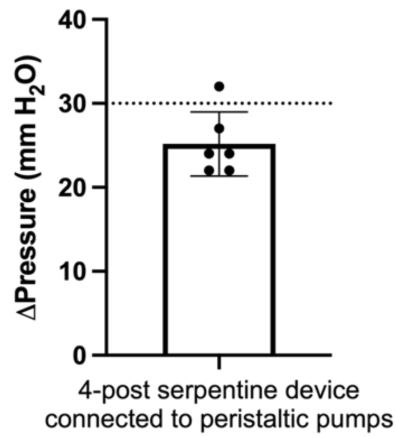


B

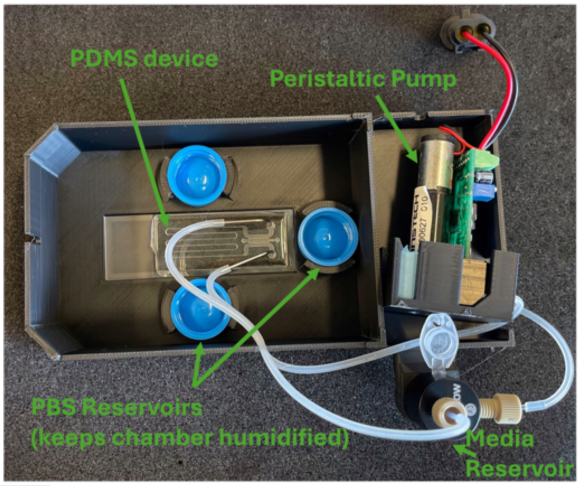


C

| | Name | Components | Sterilization |
|---|---|---|---|
| 1 | Inlet connector | • Idex® flangeless male nut, 1/16" O.D. tubing (Cole Parmer, USA)<br>• Idex® 1/16" flangeless ferrule (Cole Parmer, USA)<br>• 70mm of Idex® high-purity PFA tubing, 0.5mm I.D. 1/16" O.D. (Cole Parmer, USA) | Ethylene Oxide |
| 2 | External tubing | • 173mm of Tygon 0.031" I.D. 0.094" O.D. (VWR, USA) | Autoclave |
| 3 | Syringe tip | • 18-gauge, 90-degree angle syringe tip (McMaster Carr, USA) | Ethylene Oxide |
| 4 | External tubing | • 165mm of Tygon 0.031" I.D. 0.094" O.D. (VWR, USA) | Autoclave |
| 5 | Barbed connectors | • Idex® 1/16" Barbed Straight Connector (Cole Parmer, USA) | Autoclave |
| 6 | Pump tubing | • 63mm of Puriflex 0.44mm I.D. autoanalysis tubing (VWR, USA) | Autoclave |
| 7 | External tubing | • 67mm of Tygon 0.031" I.D. 0.094" O.D. (VWR, USA) | Autoclave |
| 8 | Outlet connector | • Idex® flangeless male nut, 1/16" O.D. tubing (Cole Parmer, USA)<br>• Idex® 1/16" flangeless ferrule (Cole Parmer, USA)<br>• 25mm of Idex® high-purity PFA tubing, 0.5mm I.D. 1/16" O.D. (Cole Parmer, USA) | Ethylene Oxide |
| X | Media Reservoir | • Microfluidic Reservoir for 1.5 mL Eppendorf® - XS (Darwin Microfluidics, France) | Ethylene Oxide |

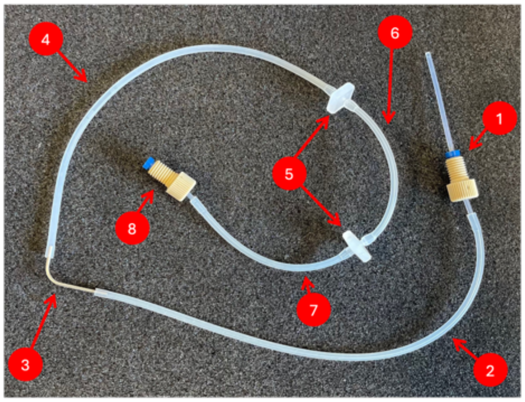


**Supplemental Figure 4: A)** Serpentine devices were measured to have a 25mm pressure drop across the channel. Each data point represents a different device and pump setup tested. B) Pumped experiment setup. PDMS devices were bonded to a glass slide and placed into a 3D printed holder connected to tubing and a peristaltic pump with the same footprint as a well plate to allow for easy imaging. C) List and image of components involved in the tubing setup and information for sterilization.

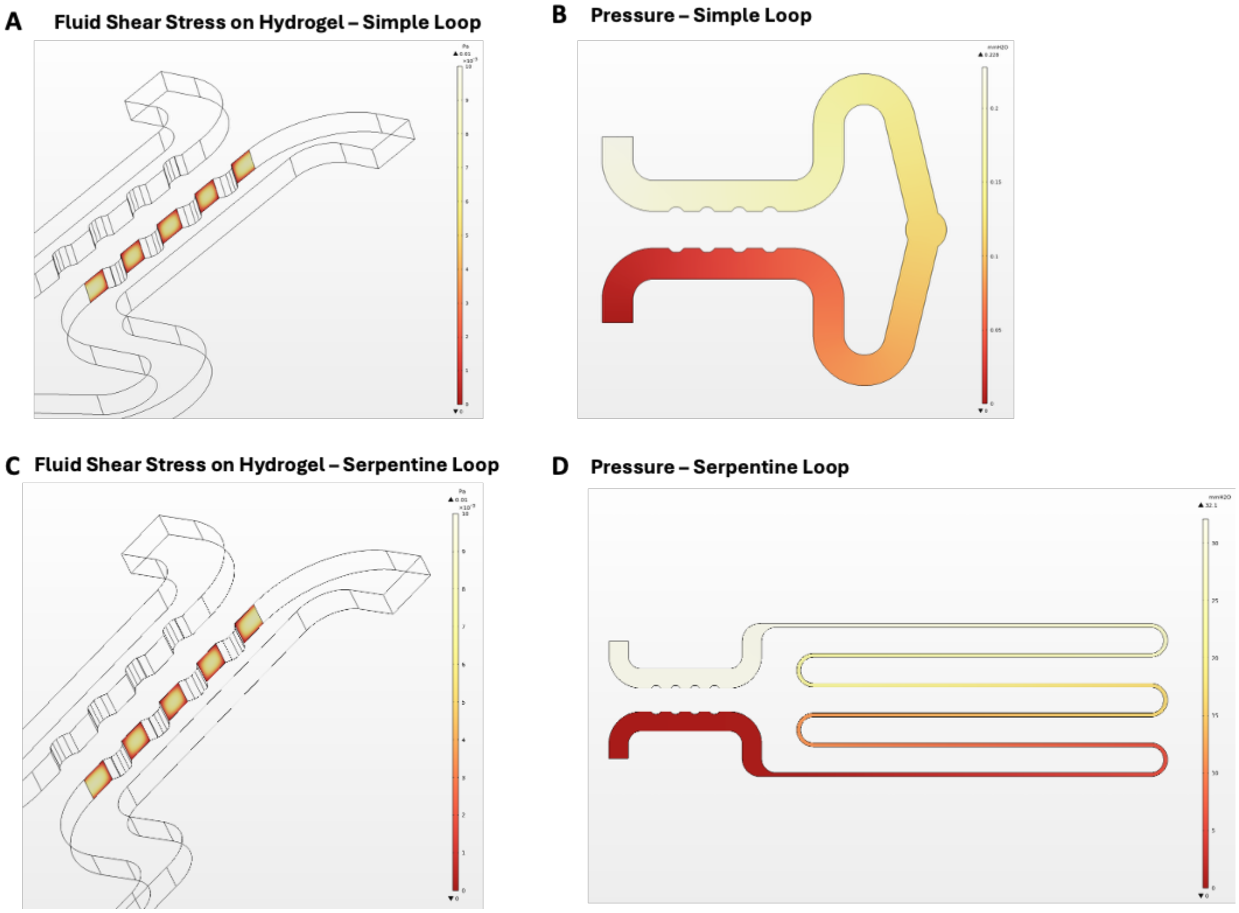


**Supplemental Figure 5:** COMSOL simulations of A) fluid shear stress on hydrogel and B) pressure in the 4-post simple-loop device used in this manuscript. COMSOL simulations of C) fluid shear stress on hydrogel and D) pressure in the 4-post serpentine-loop device used in this manuscript.

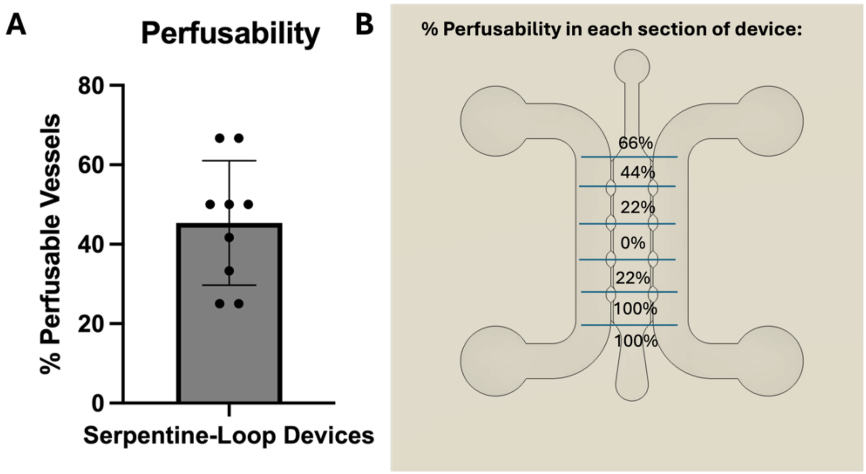


**Supplemental Figure 6:** A) Quantification of perfusable vessels throughout the entire device. Quantification of perfusability was performed at experiment end point between 7 and 10 days for 10 devices. B) Looking at each section of the device, we see all devices exhibiting perfusability in the bottom portion of the device, with most devices having perfusable vessels in the top and bottom of the device.

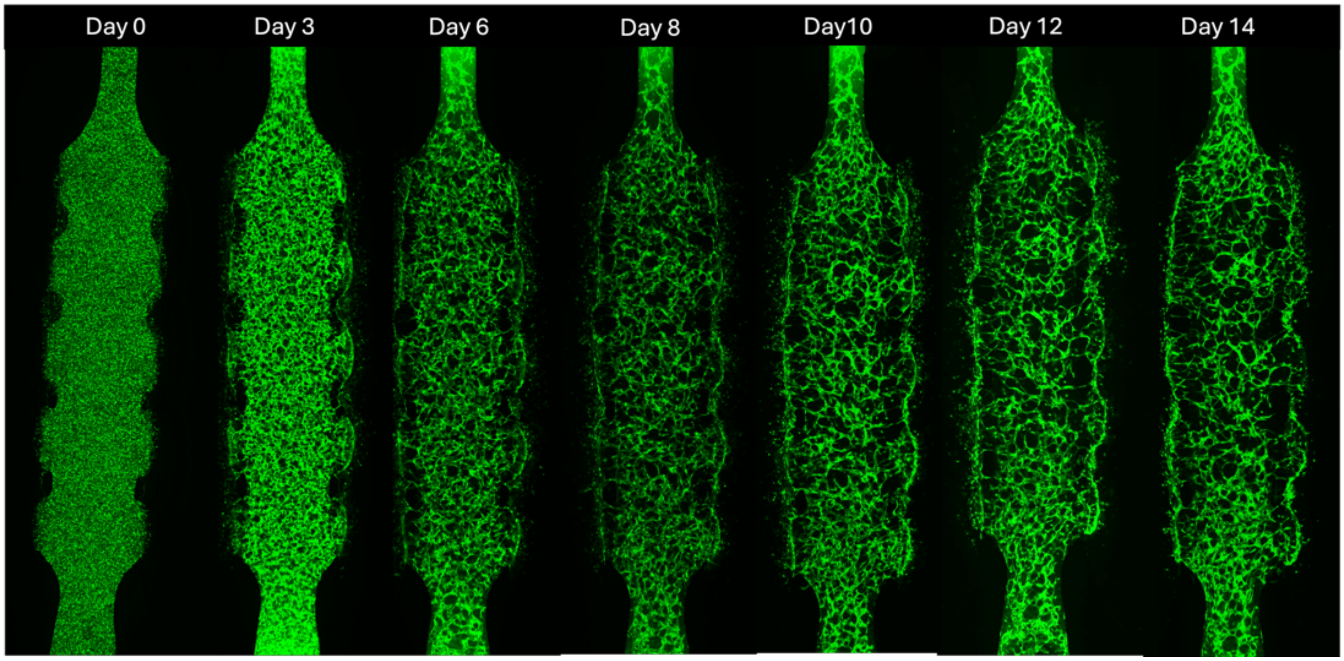


**Supplemental Figure 7:** Vessels maintain stability for at least 14 days cultured under continuous flow.

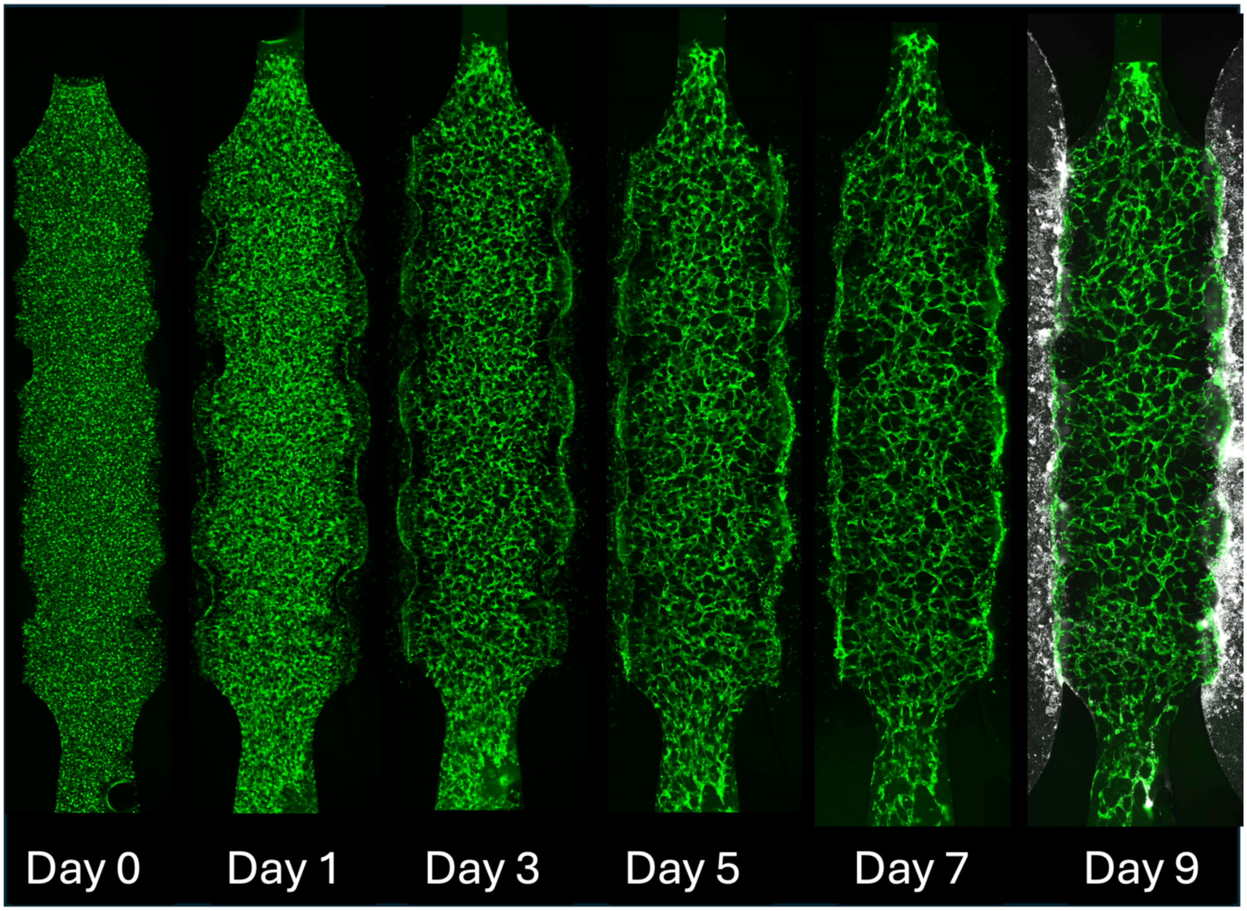


**Supplemental Figure 8:** In attempts to decrease the concentration of endothelial cells, we tested 15M HUVECs/mL with a 7:1 HUVEC:NHLF ratio. While not as successful as 18M HUVECs/mL, perfusability was observed in the bottom quarter of the device.

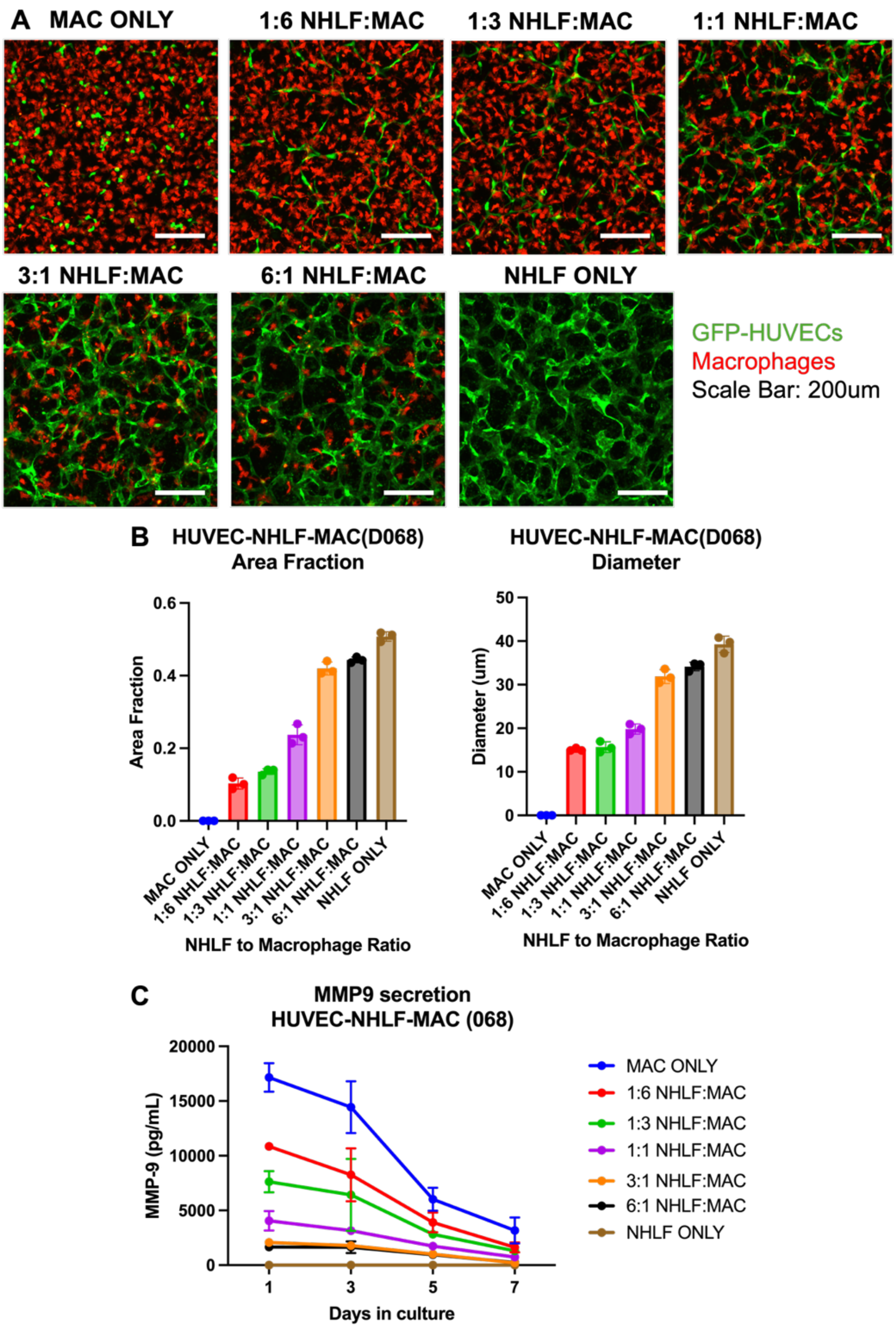


**Supplemental Figure 9: Second macrophage donor in HUVEC-NHLF-macrophage networks**. A) Day 5 static confocal images of HUVEC-NHLF-macrophage networks formed in static culture with varying ratios of macrophages to lung fibroblasts with a different macrophage donor. B) Increasing number of macrophages decreases area fraction and mean endothelial network segment diameter at day 5. C) Increasing MMP-9 secretion with increasing number of macrophages, demonstrating functionality. Data represented as mean±standard deviation. N=3.

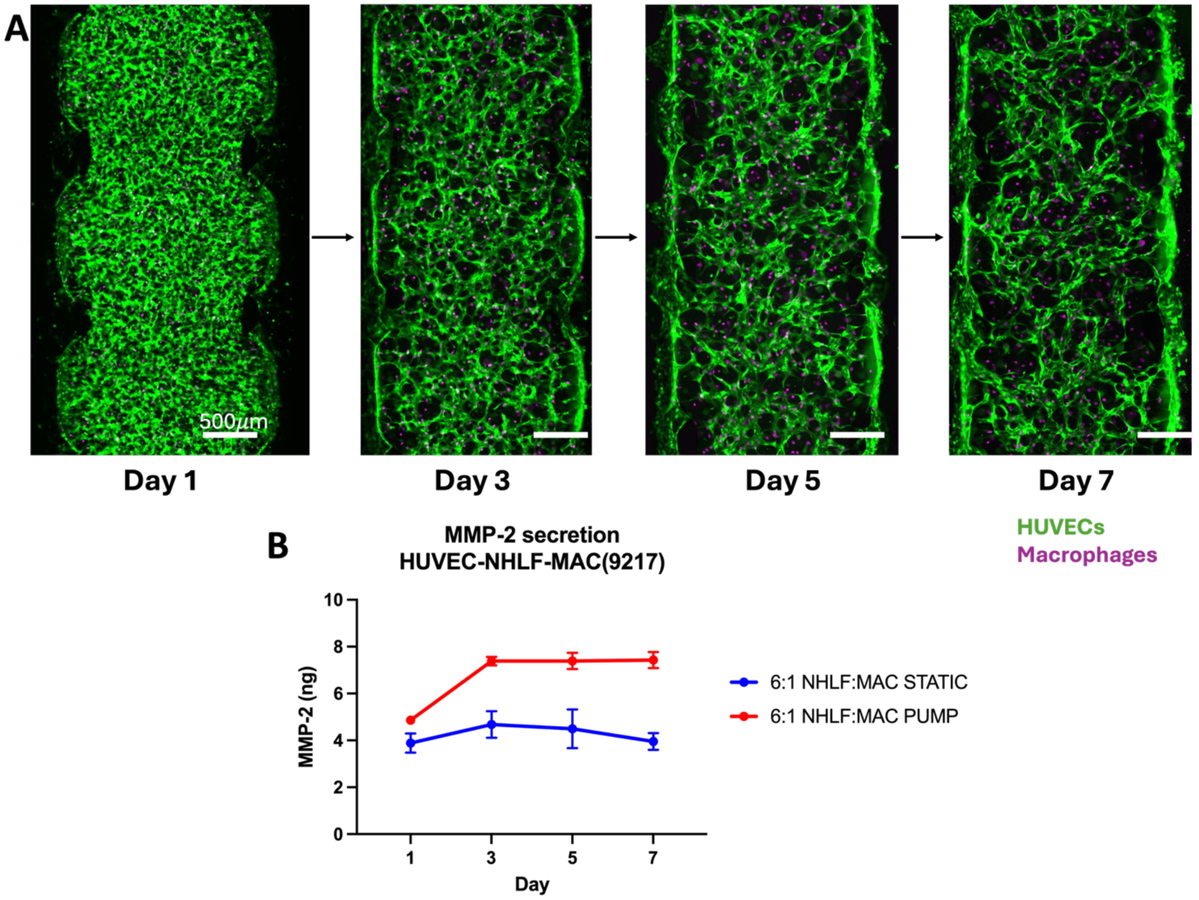


**Supplemental Figure 10: Pumped HUVEC-NHLF-macrophage networks**. A) Time course imaging of endothelial networks with a 6:1 nhlf to macrophage ratio in continuous pumped culture in serpentine devices. B) HUVEC-NHLF-macrophage networks similarly demonstrate an increase in MMP-2 detected in the supernatant compared to static culture. Data is represented as mean ± standard deviation. N=3.

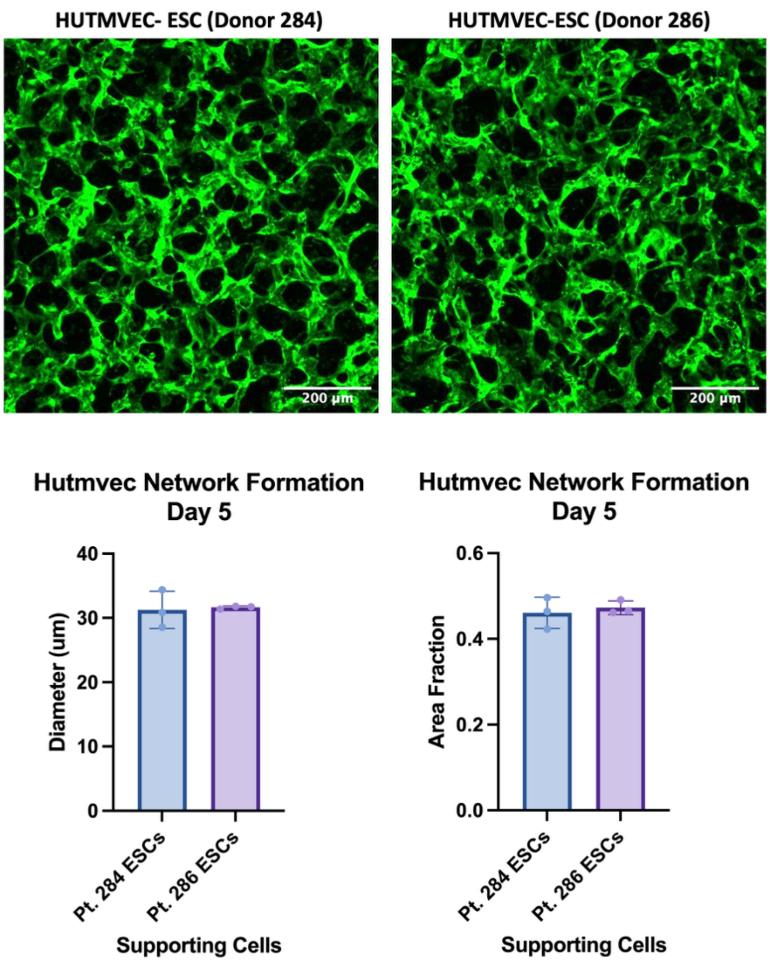


**Supplemental Figure 11:** HUTMVEC-ESC networks form similarly with two different endometrial stromal cell donors with no significant difference in area fraction or diameter in static culture. Data is represented as mean ± standard deviation. N=3.

A

| | |
|---|---|
| Vasculife Basal Medium | 50% base |
| DMEM/F12 | 50% base |
| FGF basic | 2.5ng/mL |
| Ascorbic Acid | 25 µg/mL |
| Hydrocortisone Hemisuccinate | 0.5 µg/mL |
| C/S FBS | 2.5% |
| L-Glutamine | 5 mM |
| rhIGF-1 | 7.5ng/mL |
| rhEGF | 27.5 ng/mL |
| rhVEGF | 2.5 ng/mL |
| Heparin Sulfate | 0.375 U/mL |
| Gentamicin | 15 mg/mL |
| Amphotericin B | 7.5 µg/mL |
| B27 Supplement | 0.5x |
| N2 supplement | 0.5x |
| PenStrep | 0.5% |
| Glutamax | 0.5x |
| ITS | 0.5% |
| rhNoggin | 50 ng/mL |
| Nicotinamide | 0.5 mM |
| N-Acetyl-L-Cysteine | 0.625 mM |
| rhFGF-10 | 25 ng/mL |
| rhRspondin-1 | 100 ng/mL |
| Estradiol | 10nM |

B

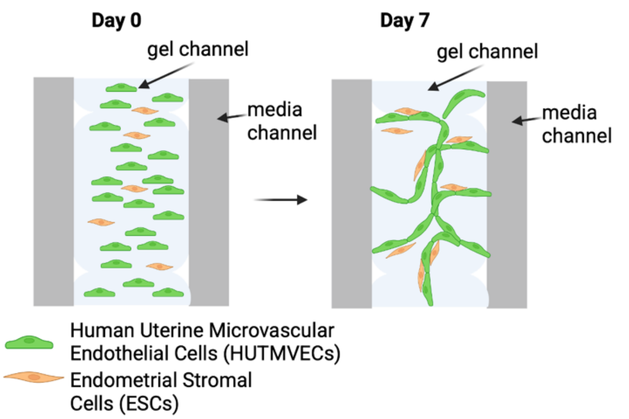


C

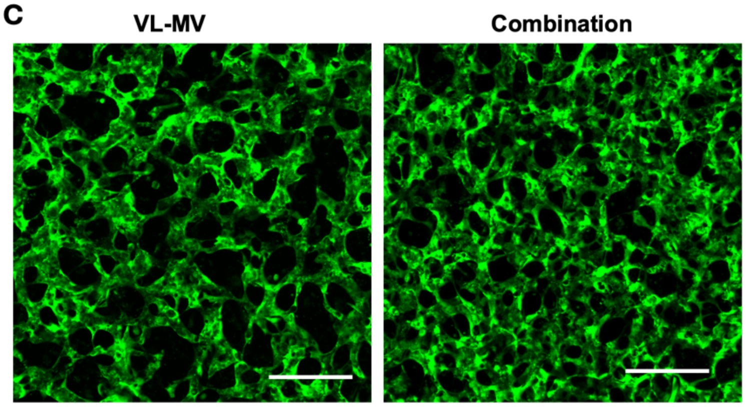


D

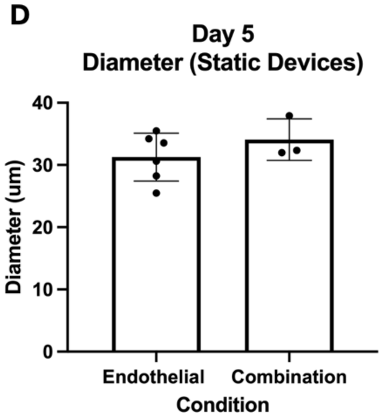


**Supplemental Figure 12: Testing combination media.** A) Composition of combination media. B) Using HUTMVEC-ESC culture, we tested a combination media (50% microvascular endothelial media, 50% endometrial organoid media without A8301 and Y27632) to form microvascular networks in static culture. C)Networks formed to a similar extent in the combination media D) with no significant difference in network diameters. Scale Bar: 200µm.

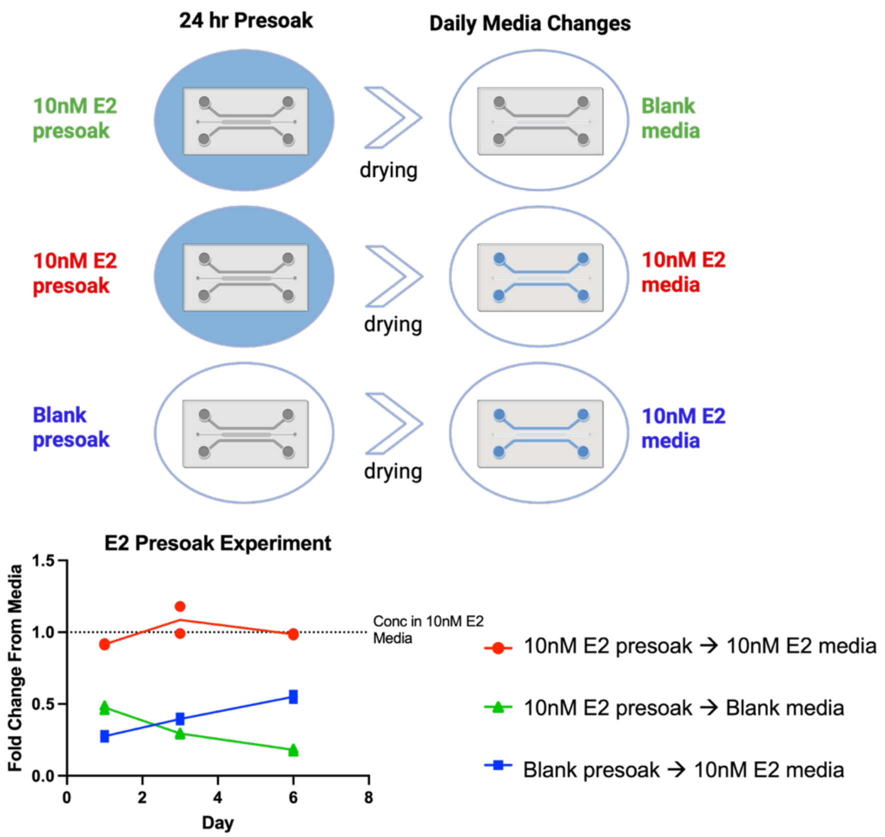


**Supplementary Figure 13: E2 Presoak test.** For a final experiment to combine endometrial epithelial organoids and vessels, it was necessary to use an estradiol containing medium to support organoid growth. As PDMS rapidly absorbs lipophilic compounds, it was necessary to develop a method to pre-saturate the PDMS to maintain a concentration of 10nM E2 throughout the duration of the experiment. Presoaking the devices for 24 hours in a 10nM E2 containing solution, prior to loading the gel maintained the concentration throughout the experiment by using media containing 10nM E2. If devices were not presoaked, the media only contained approximately 50% of the desired concentration by day 6. N=2 devices per condition.
